# Multi-omic analysis of medium-chain fatty acid synthesis by *Candidatus* Weimerbacter bifidus, gen. nov., sp. nov., and *Candidatus* Pseudoramibacter fermentans, sp. nov.

**DOI:** 10.1101/726943

**Authors:** Matthew J. Scarborough, Kevin S. Myers, Timothy J. Donohue, Daniel R. Noguera

## Abstract

Chain elongation is emerging as a bioprocess to produce valuable medium-chain fatty acids (MCFA; 6 to 8 carbons in length) from organic waste streams by harnessing the metabolism of anaerobic microbiomes. Although our understanding of chain elongation physiology is still evolving, the reverse β-oxidation pathway has been identified as a key metabolic function to elongate the intermediate products of fermentation to MCFA. Here, we describe two chain-elongating microorganisms that were enriched in an anaerobic microbiome transforming the residues from a lignocellulosic biorefining process to short- and medium-chain fatty acids. Based on a multi-omic analysis of this microbiome, we predict that *Candidatus* Weimerbacter bifidus, gen. nov., sp. nov. used xylose to produce MCFA, whereas *Candidatus* Pseudoramibacter fermentans, sp. nov., used glycerol and lactate as substrates for chain elongation. Both organisms are predicted to use an energy conserving hydrogenase to improve the overall bioenergetics of MCFA production.

**IMPORTANCE:** Microbiomes are vital to human health, agriculture, environmental processes, and are receiving attention as biological catalysts for production of renewable industrial chemicals. Chain elongation by MCFA-producing microbiomes offer an opportunity to produce valuable chemicals from organic streams that otherwise would be considered waste. However, the physiology and energetics of chain elongation is only beginning to be studied, and we are analyzing MCFA production by self-assembled communities to complement the knowledge that has been acquired from pure culture studies. Through a multi-omic analysis of an MCFA-producing microbiome, we characterized metabolic functions of two chain elongating bacteria and predict previously unreported features of this process.

## INTRODUCTION

Chain elongation has been proposed as a microbial process to produce valuable chemicals from complex waste feedstocks (1). This bioprocess relies on the combined metabolism of an anaerobic microbiome to hydrolyze complex organic substrates, ferment the hydrolyzed products to small organic intermediates (2 to 3 carbon molecules), and elongate these fermentation products to medium-chain fatty acids (MCFA; 6 to 8 carbon molecules) through reverse β-oxidation (1). MCFA are an attractive product due to their high value, relatively low solubility in water, and potential to offset fossil fuel demands for petrochemicals and other products (2, 3). Bioreactors performing chain elongation also provide model systems for studying the metabolic contributions of uncultured organisms to this process. To date, all but one of the known chain-elongating bacteria belong to the *Clostridia* class within the *Firmicutes* phylum (1). These chain-elongating bacteria primarily use lactate (4, 5), ethanol (6), or carbohydrates (7) to drive MCFA production.

We recently described a chain-elongating microbiome that produced sufficient hexanoic and octanoic acids from lignocellulosic biorefining residues to reduce the minimum selling price of ethanol produced in a biorefinery (2). Using a combination of metagenomics and metatranscriptomics, we characterized this microbiome as having a small set of high abundance organisms (8), with two populations within the *Clostridia* class performing chain elongation. One high abundance member of this microbiome (LCO1) (8) belonged to the *Lachnospiraceae* family and was predicted to produce MCFA from xylose and other pentoses; the second one (EUB1) (8) corresponded to the *Eubacteriaceae* family and was predicted to produce MCFA from lactate.

Here, we combined multi-omic approaches to further analyze the genomic and metabolic features of these two predicted MCFA-producing organisms. A time-series gene expression analysis showed that transcripts encoding proteins predicted to be involved in reverse β-oxidation are among the most abundant transcript RNAs after feeding the lignocellulosic biorefinery residues to the microbial community. Our analysis also reveals that both organisms contain transcripts that encode a proton-translocating energy conserving hydrogenase, suggesting contributions of previously unreported metabolic networks to MCFA production. Based on these new results, we conclude that LCO1 represents a novel genus within the *Lachnospiraceae* family and propose the name of *Candidatus* Weimerbacter bifidus, gen. nov., sp. nov. Our data also predicts that EUB1 represents a new species within the *Pseudoramibacter* genus, and we propose the name *Candidatus* Pseudoramibacter fermentans sp. nov., to represent this new species.

## RESULTS AND DISCUSSION

### Refinement of metagenome-assembled genomes (MAGs)

We previously reported the construction of draft MAGs from a MCFA-producing microbiome fed with lignocellulosic biorefinery residues, in which LCO1 and EUB1 represented the abundant chain-elongating microorganisms (8). These draft MAGs were constructed using DNA samples from the first 120 days of reactor operation. To improve the quality of these MAGs, we obtained Illumina and PacBio sequencing reads obtained from the same microbiome at different times during a 378-day operational period. We co-assembled 244 million Illumina Hi-seq (2×250) reads from five time points (days 96, 120, 168, 252, and 378; **Fig. S1**) into 24,000 contigs. Contigs were binned into MAGs; the MAGs with relative abundance greater than 1% were then gap-filled with PacBio reads from the day 378 sample. This analysis resulted in an overall improvement in MAG quality with respect to completeness, contamination, and number of scaffolds (**Table 1**). The two most abundant MAGs derived from this analysis, LCO1.1 and EUB1.1 (accounting for ~80% of the recovered DNA sequences; **Supplementary Data File 1**), provide improved predictions for the genetic makeup of the organisms previously denoted as LCO1 and EUB1 (8), respectively.

**Table 1.** Summary of metagenome-assembled genomes from MCFA-producing bioreactor

| MAG Name | Original Name<sup>1</sup> | Relative Abundance<sup>2</sup> (%) | Completeness (%)<sup>3</sup> | Contamination (%)<sup>3</sup> | Genome size (Mbp)<sup>3</sup> | No. scaffolds<sup>3</sup> |
| --- | --- | --- | --- | --- | --- | --- |
| *Cand. W. bifidus* (LCO1.1) | LCO1 | 75.3 | 96.9 [95.4] | 0.5 [0.0] | 2.39 [2.10] | 10 [44] |
| *Cand. P. fermentans* (EUB1.1) | EUB1 | 4.7 | 99.2 [97.8] | 0.2 [0.2] | 2.29 [2.00] | 29 [35] |
| COR1.1 | COR1 | 2.4 | 95.0 [99.2] | 6.7 [0.8] | 2.41 [2.51] | 82 [225] |
| COR3.1 | COR3 | 2.8 | 98.4 [98.4] | 2.4 [7.4] | 3.02 [3.65] | 134 [533] |
| COR4.1<sup>4</sup> |  | 1.1 | 100 | 0.7 | 2.45 | 8 |
| LAC1.1 | LAC1 | 3.8 | 99.5 [99.5] | 1.1 [1.1] | 2.77 [2.63] | 9 [18] |
| LAC2.1 | LAC2 | 2.0 | 99.4 [99.4] | 1.6 [1.6] | 3.18 [3.18] | 37 [79] |
| LAC4.1 | LAC4 | 1.6 | 97.7 [98.9] | 0.6 [1.3] | 3.14 [3.35] | 53 [95] |
| LAC5.1 | LAC5 | 2.5 | 99.2 [80.1] | 0.0 [0.8] | 2.11 [1.48] | 6 [181] |
| LAC6.1<sup>4</sup> |  | 1.9 | 99.1 | 1.1 | 2.80 | 12 |
| LAC7.1<sup>4</sup> |  | 2.0 | 99.1 | 2.8 | 3.41 | 33 |

<sup>1</sup> Names reported in Scarborough et al. (8)
<sup>2</sup> Relative abundance based in DNA read mapping normalized to genome size for day 252 sample.
<sup>3</sup> Numbers in brackets indicate values for assembled MAGs reported in Scarborough et al. (8)
<sup>4</sup> MAGs not assembled in Scarborough et al. (8)

### Phylogenetic classification of LCO1.1 and EUB1.1

The Genome Taxonomy Database (GTDB) Tool kit (9) was used to provide a taxonomic classification of LCO1.1 and EUB1.1 (**Fig. 1**). The reconstructed LCO1.1 genome, which accounted for ~75% of the mapped DNA reads (**Table 1**), clusters with a group of genomes identified in the GTDB with the name UBA2727 within the *Lachnospiraceae* family, in the order *Clostridia*, phylum *Firmicutes*. This cluster contains three MAGs obtained from rumen samples (GCA_002474355.1, GCA_002480925.1, GCA_900316165.1) (9), as well as a genome of an isolate from the Hungate 1000 project (named as *Lachnospiraceae* bacterium C10 in the NCBI database; GCF_900100095.1) (10) (**Supplementary Data File 2**). A comparative analysis of LCO1.1 with the four representatives of the UBA2727 cluster (**Fig. 1B**) shows average nucleotide identity (ANI) values greater than 70% with the three MAGs and 68% with the Hungate 1000 project isolate. The most closely-related type strain is *Shuttleworthia satelles*, an organism isolated from the human oral cavity and described as a carbohydrate consumer and butyrate producer (11). Given the low ANI value (66%) obtained when the LCO1.1 genome is compared to that of *S. satelles*, we propose here the name *Candidatus* “Weimerbacter” to define the organisms comprised of the UBA2727 cluster plus the LCO1.1 MAG. Further, we propose the epithet “bifidus” to describe a species within this new genus, represented by LCO1.1.

**Figure 1.**
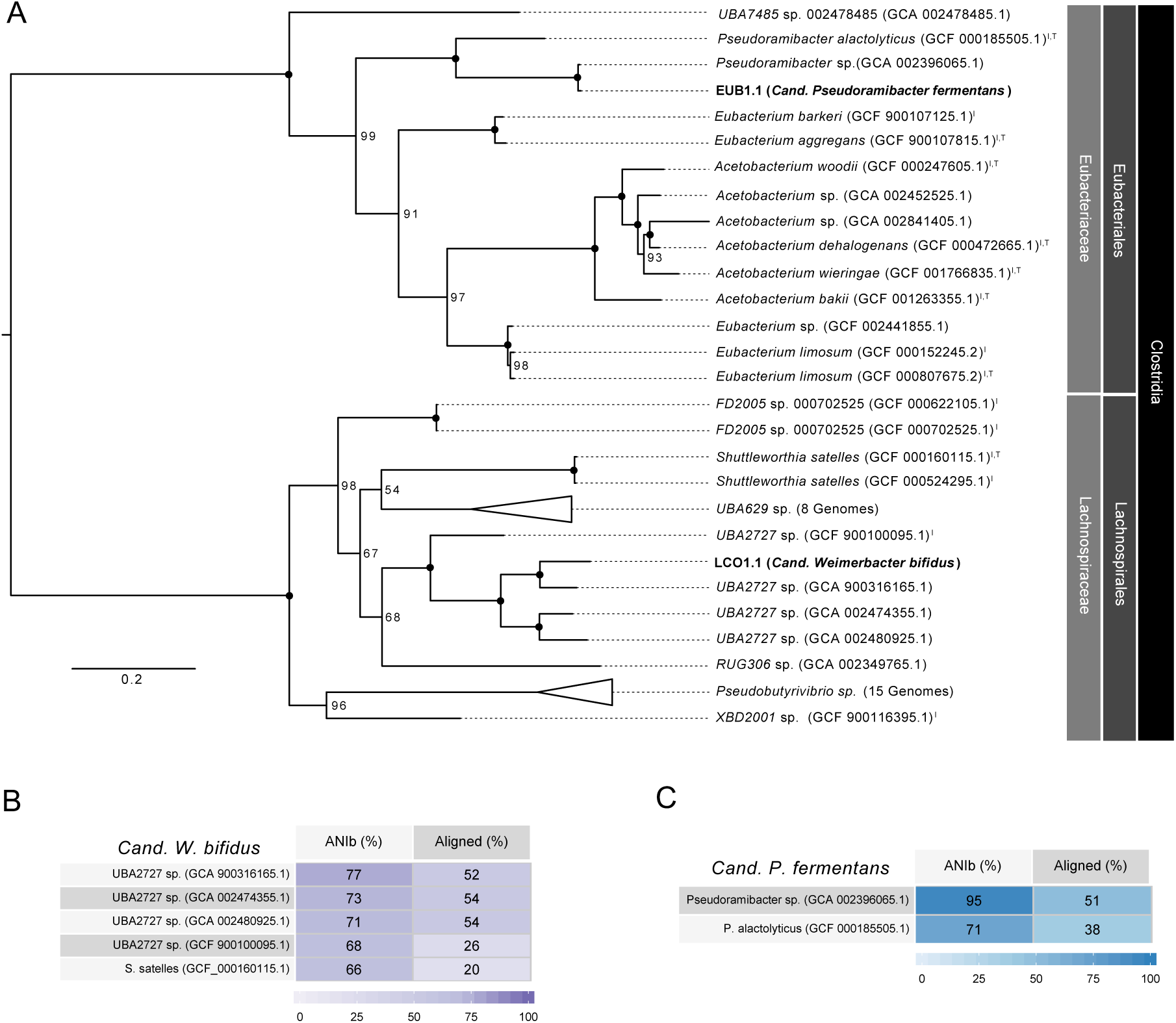
Phylogenetic analysis of two metagenome-assembled metagenomes (shown in bold text) predicted to perform chain elongation in an anaerobic microbiome converting lignocellulosic residues to medium-chain fatty acids. (**A**) A phylogenetic tree was created based on concatenated amino acid sequences for 120 single copy marker genes from the GTDB using RAxML. Bootstrap support values are indicated at nodes. Closed circles indicate bootstrap support values of 100. Organisms that have representative isolates are shown indicated by an “I” superscript and organisms representing NCBI type strains are indicated with a “T” superscript. Genome identifiers are provided in parentheses; (B) BLAST Average Nucleotide Identity (ANIb) comparison for LCO1.1 (*Cand*. W. bifidus) to related genomes; (C) ANIb comparison for EUB1.1 (*Cand*. P. fermentans) to related genomes.

The EUB1.1 MAG, which accounted for ~5% of the mapped DNA reads (**Table 1**), clustered in the *Pseudoramibacter* genus, within the *Eubacteriaceae* family, also in the order *Clostridia*. This genus currently contains only one known species, defined by the isolate *P. alactolyticus* (originally named *Ramibacterium alactolyticum* (12)), which was recovered from a human lung abscess and has been shown to produce short- and medium-chain fatty acids (12). The ANI of EUB1.1 to *P. alactolyticus* is 71%, with an alignment factor of 38% (**Fig. 1C**). The GTDB also contained a related MAG that was obtained from a sheep rumen metagenomic analysis (9), and identified as *Pseudoramibacter* sp. (GCA 002396065.1). This MAG had an ANI of 95% and an alignment factor of 51% to EUB1.1 (**Fig. 1C**). Its completion was estimated to be only 57%, which likely contributes to its low alignment factor to EUB1.1. A comparative analysis of these three *Pseudoramibacter* genomes (**Supplementary Data File 3**) shows that among all similarities, they all contain genes typically associated with reverse β-oxidation, consistent with their predicted ability to produce MCFA. Based on the clustering of EUB1.1 within the *Pseudoramibacter* genus, on the obtained ANI values, and on the comparative analysis that follows, we propose here a new species within the *Pseudoramibacter* genus, represented by EUB1.1 and *Pseudoramibacter* sp., and identified with the epithet “fermentans.”

### Multi-omic analysis of *Cand*. W. bifidus and *Cand*. P. fermentans

To evaluate the potential of *Cand*. W. bifidus and *Cand*. P. fermentans to synthesize MCFA, we analyzed transcript abundance as a function of time after the bioreactor received a transient load of lignocellulosic biorefinery residue (**Supplementary Data Files 4 and 5**). During this experiment, we collected eleven samples for transcriptomic analysis over a 36-hour time period. Concurrent analysis of the media over this time period showed that xylose and glycerol were consumed whereas lactate transiently accumulated in the reactor (**Fig. S2**). We used this multi-omic analysis to investigate substrate utilization, the enzymes involved in converting substrates into intermediates and end-products, the potential for MCFA production via the reverse β-oxidation pathway, and the predicted energy conserving features of the predominant MAGs in this anaerobic microbiome.

#### Chain elongation in *Cand*. W. bifidus

Four enzymes are known or predicted to comprise the reverse β-oxidation cycle (**Fig. 2**). In the first step, acyl-CoA acetyl transferase (ACAT) condenses an acetyl-CoA with an acyl-CoA; the product of this reaction is reduced by 3-hydroxy-acyl-CoA dehydrogenase (HAD), followed by a dehydration by 2-enoyl-CoA with enoyl-CoA dehydratase (ECoAH) and a reduction by an acyl-CoA dehydrogenase (ACD) to form an elongated acyl-CoA. In some organisms, the last dehydrogenation reaction is catalyzed by an electron-bifurcating energy-conserving enzyme where the enoyl-CoA reduction with NADH is paired with the reduction of ferredoxin through an ACD complex containing the electron transfer flavoproteins 7 EtfA and EtfB (13). The *Cand*. W. bifidus genome has a gene cluster encoding ACAT, HAD, ACD, EtfA, and EtfB (**Fig. 3A**), while a gene predicted to encode ECoAH is located on a different region of the genome. The abundance of transcripts encoding reverse β-oxidation enzymes was at or above the 90^th^ percentile in all samples analyzed (**Fig. 4A**). Pairwise comparisons of reverse β-oxidation transcript abundance show high correlations for genes in the cluster encoding ACAT, HAD, ACD, EtfA, and EtfB over the course of this analysis, but low correlations with the ECoAH transcript (**Fig. S3**) suggesting that this later gene is not co-expressed with those predicted to be involved in the reverse β-oxidation pathway of *Cand*. W. bifidus.

**Figure 2.**
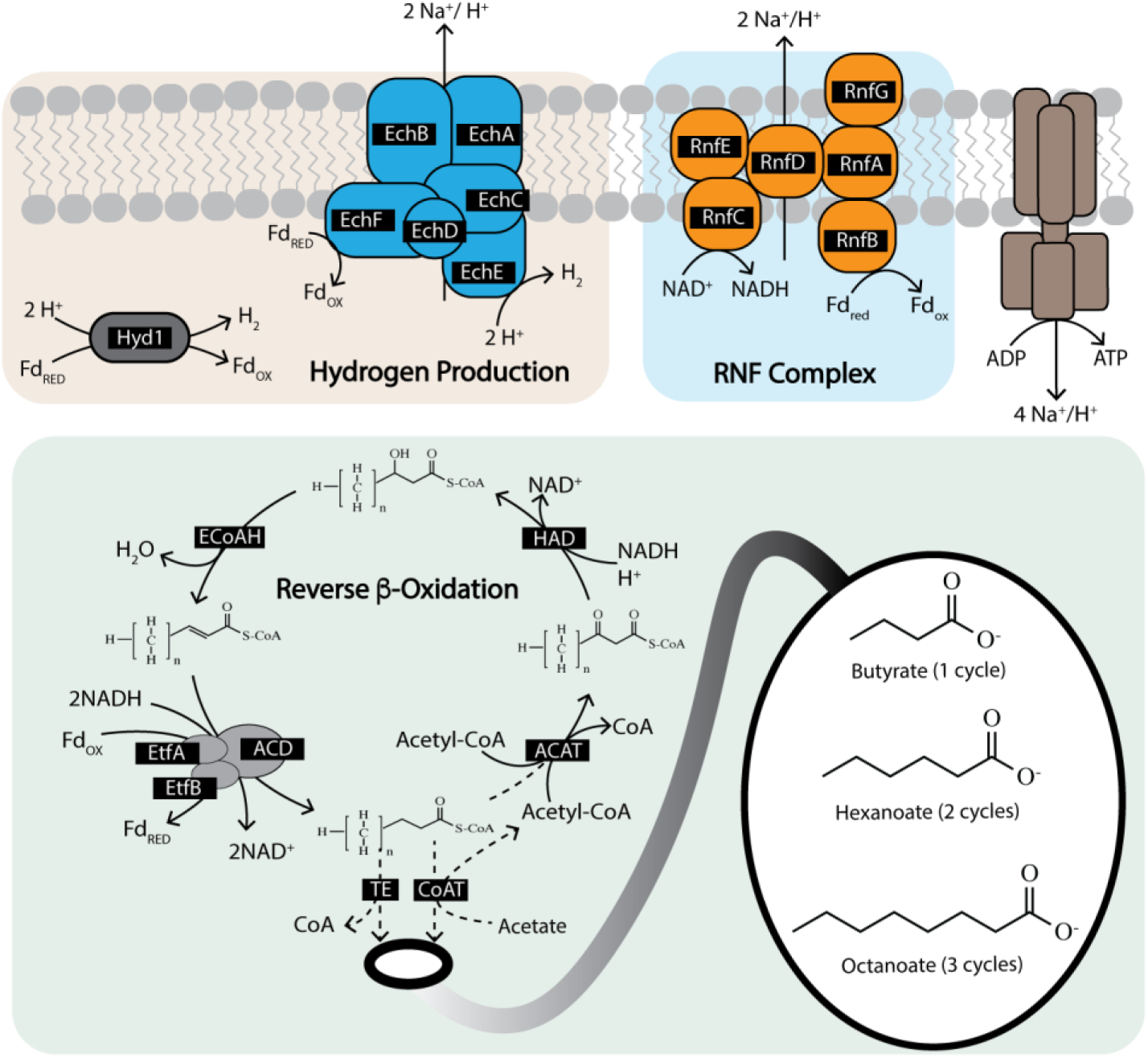
Metabolic pathways involved in chain elongation. Reverse β-oxidation is a four-step process using acyl-CoA acetyl transferase (ACAT), 3-hydroxy-acyl-CoA dehydrogenase (HAD), enoyl-CoA dehydratase (ECoAH), and acyl-CoA dehydrogenase (ACD). The reduction of enoyl-CoA with NADH can be combined with the reduction of ferredoxin through an electron-bifurcating acyl-CoA dehydrogenase complex containing EtfA and EtfB. The terminal enzyme of reverse β-oxidation can be either a CoA transferase (CoAT) or thioesterase (TE). Reverse β-oxidation is coupled with proton-translocating enzymes to generate ATP with reduced ferredoxin.

**Figure 3.**
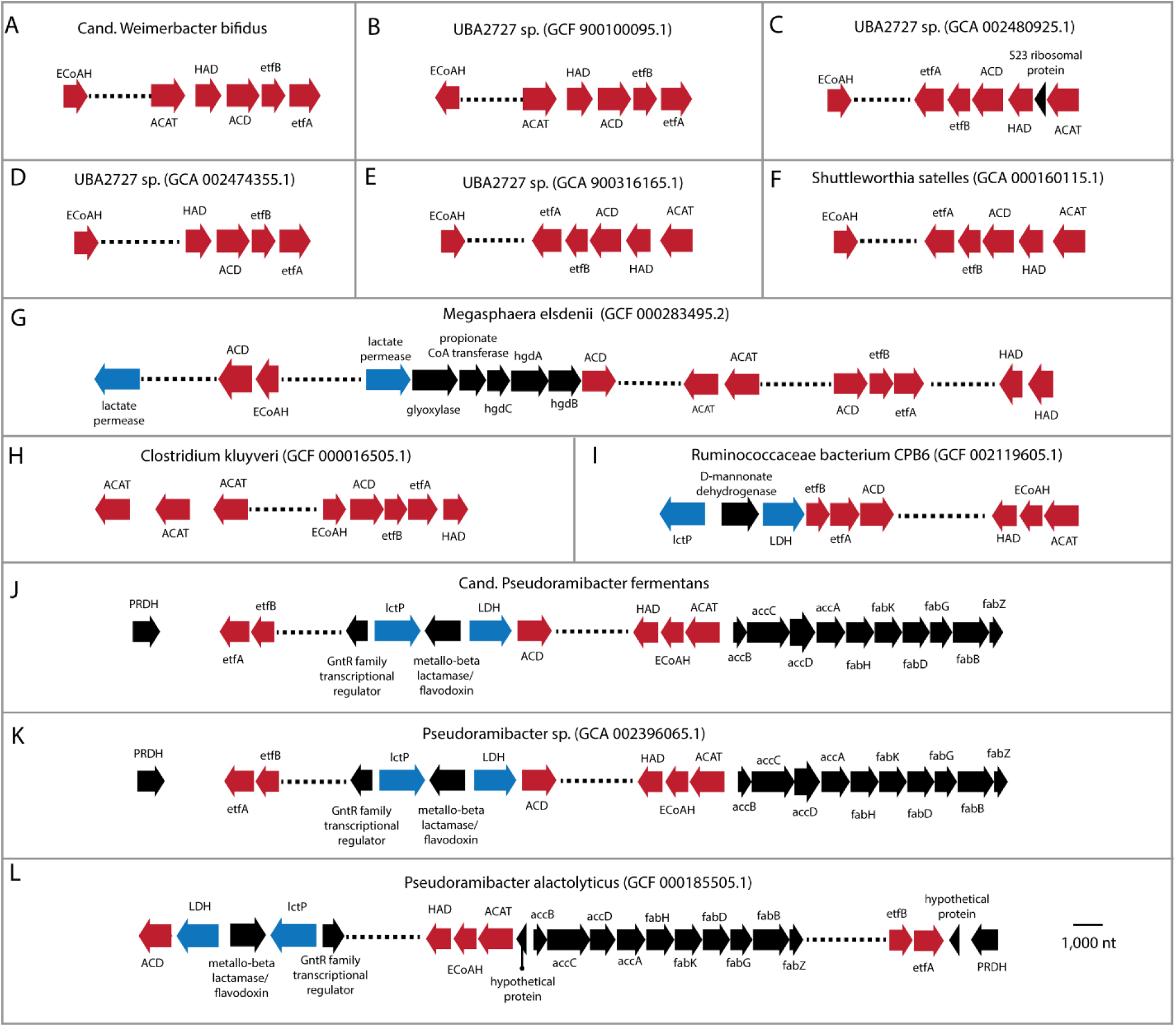
Organization of selected genes in organisms related to *Cand*. W. bifidus*, Cand*. P. fermentans, and known chain-elongating organisms. Reverse β-oxidation genes are shown in red and genes involved in lactate utilization, if present, are shown in blue. Reverse β-oxidation genes include acyl-CoA acetyl transferase (ACAT), 3-hydroxy-acyl-CoA dehydrogenase (HAD), enoyl-CoA dehydratase (ECoAH), acyl-CoA dehydrogenase (ACD), electron transfer flavoprotein A (*etfA*), and electron transfer flavoprotein B (*etfB*). Lactate utilization genes include lactate permease (*lctP*) and lactate dehydrogenase (LDH). Other genes include 2-hydroxyglutaryl-CoA dehydratase (*hgd*) involved in propionate production in *Megasphaera elsdenii* (G) that are adjacent to *lctP* and ACD; prephenate dehydrogenase (PRDH); and fatty acid biosynthesis genes when they are adjacent to reverse β-oxidation genes including acetyl-CoA carboxylase (*acc*) and fatty acid biosynthesis genes (J,K,L).

**Figure 4.**
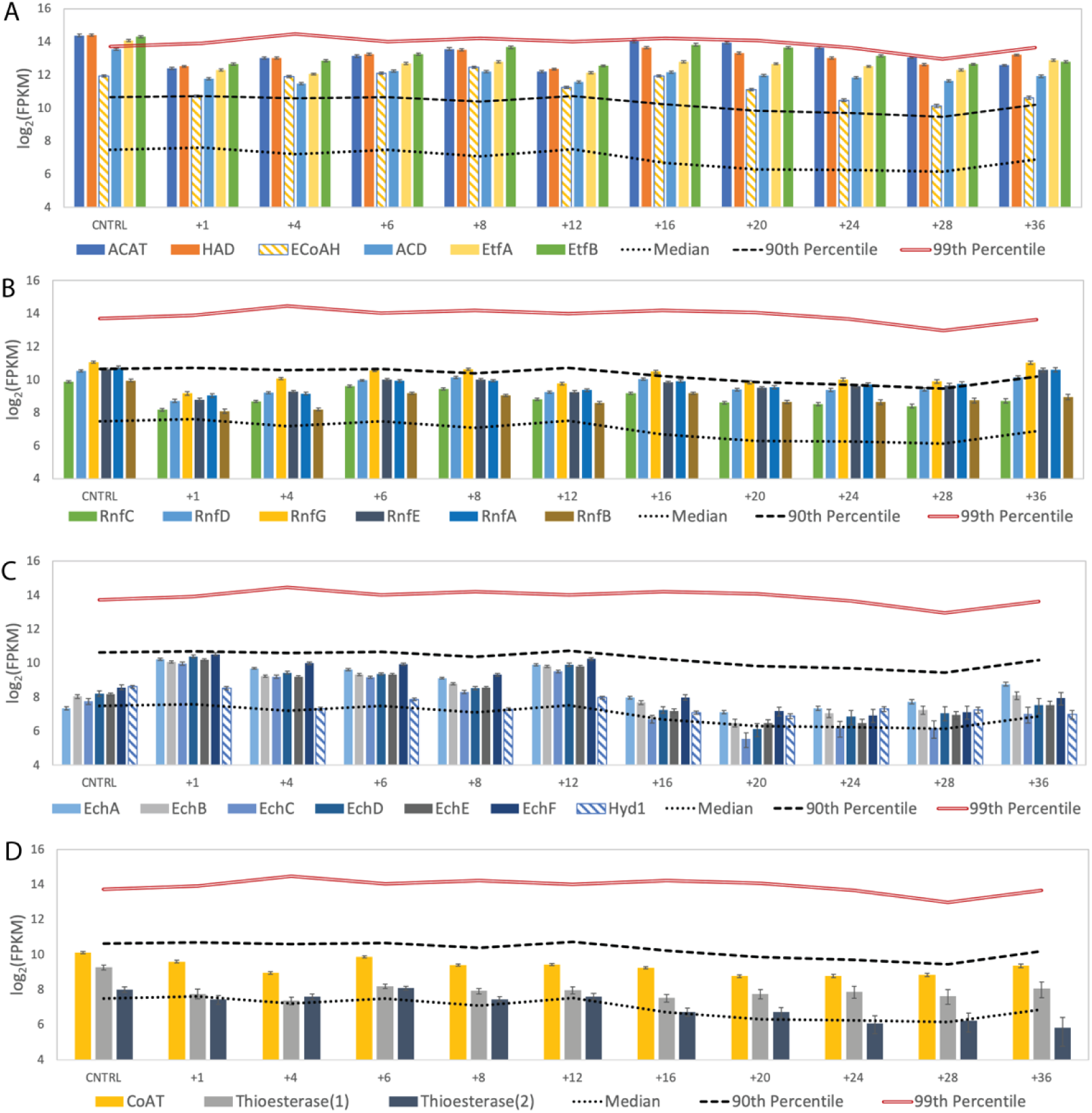
Expression of predicted chain elongation genes by *Cand*. W. bifidus. Transcript abundance data is shown for transcripts encoding enzymes used for (A) reverse β-oxidation, (B) the RNF complex, (C) hydrogenases, and (D) terminal enzymes of reverse β-oxidation. The roles of the designated enzymes in chain elongation are provided in Fig. 2.

Producing reduced ferredoxin during reverse β-oxidation constitutes a potential energy conserving process (**Fig. 2**) because its subsequent oxidation can be used to create an ion motive force by either the RNF complex (RnfABCDEG), as proposed for the MCFA-producing *C. kluyveri* (14), or the energy conserving hydrogenase complex Ech, as described for *C. thermocellum* (15). In *Cand*. W. bifidus, the genes encoding the subunits of the RNF complex are found in a single operon arranged as *rnfCDGEAB*, which is consistent with the gene organization in *C. kluyveri*, *C. difficile*, and other *Clostridia* (16). Genes encoding individual subunits of the Ech complex are also found in a single operon in *Cand*. W. bifidus (*echABCDEF*; **Supplementary Data File 2**) and are adjacent to genes encoding HypB, HypC, and HypD, which are involved in hydrogenase assembly (17, 18). Predicted *echABCDEF* gene clusters were not found in genomes of the other known chain-elongating organisms analyzed in this study (**Supplementary Data File 6**), but they have been found in a diverse range of other bacteria and archaea (19).

Time-series transcriptomic analysis showed that *Cand*. W. bifidus produced high levels of transcripts encoding the RNF and the Ech complexes when MCFA were produced after addition of lignocellulosic biorefinery residues (**Fig. 4**). Transcripts encoding subunits of the RNF complex (RnfABCDEFG) were present above the 90^th^ percentile at several time points after addition of lignocellulosic biorefinery residues and found to be at or above median expression levels throughout the study (**Fig. 4B**). Transcripts for genes encoding subunits of the Ech hydrogenase complex follow a different pattern, with high level abundance only for the first 12 hours after lignocellulosic biorefinery residues addition (**Fig. 4C**). We also found that transcripts of Ech complex genes were more abundant than those encoding a putative periplasmic ferredoxin hydrogenase (Hyd1), the only other hydrogenase predicted to be present in the *Cand*. W. bifidus genome. Thus, the multi-omic data supports a role for both the RNF and Ech complexes during MCFA production, likely by conserving energy via generation of an ion motive force (**Fig. 2**).

The reverse β-oxidation cycle is also predicted to require either a CoA transferase (CoAT) or a thioesterase to remove the CoA from the terminal acyl-CoA molecule, thereby releasing the corresponding acid (**Fig. 2**). During the course of this experiment, *Cand*. W. bifidus expressed genes encoding one predicted CoAT and two predicted thioesterases. Transcripts of all three genes were at or above median levels throughout the time course of this experiment (**Fig. 4D**). The genes encoding putative thioesterase enzymes and CoAT are not located near other predicted reverse β-oxidation genes in the *Cand*. W. bifidus genome (**Supplementary Data File 2**), but the abundance of transcripts encoding one of the thioesterase enzymes (TE.1; **Fig. S3**) is more closely correlated with other reverse β-oxidation pathway transcripts than the other thioesterase (TE.2; **Fig. S3**). TE.1 was annotated as a thioesterase superfamily protein (**Supplementary Data File 2**), and a Basic Local Alignment Search Tool (BLAST) analysis of amino acid sequences (**Supplementary Data File 7**) places it in the hotdog-fold family of thioesterase proteins, some of which have been shown to cleave CoA from medium- and long-chain acyl-CoA molecules (20). Thus, this analysis indicates that CoAT and TE.1 may participate in the terminal step of reverse β-oxidation in *Cand*. W. bifidus.

#### *Cand*. W. bifidus is predicted to use multiple routes to consume xylose as a source of carbon for MCFA production

Previous studies (8) and the revised genome provided in this work predict that *Cand*. W. bifidus can metabolize xylose and other pentoses (**Fig. 5A**) as the main source of energy and carbon when growing in the MCFA-producing microbiome (8). In this time-series experiment, we found that transcripts encoding an ABC transporter predicted to transport multiple sugars were present at or above the 99^th^ percentile at several time points, (**Fig. 5B**) suggesting a role for this protein in sugar uptake. To investigate potential routes for sugar utilization by *Cand*. W. bifidus, we compared transcript abundance of genes encoding enzymes predicted to function in the pentose phosphate and phosphoketolase pathways after the addition of lignocellulosic biorefinery residues (**Fig. 5B**). Patterns of transcript abundance indicate that genes encoding enzymes for both pathways are expressed above median expression levels, suggesting that both pathways are used for pentose utilization and its subsequent conversion to intermediates that then enter the reverse β-oxidation pathway to produce MCFA.

**Figure 5.**
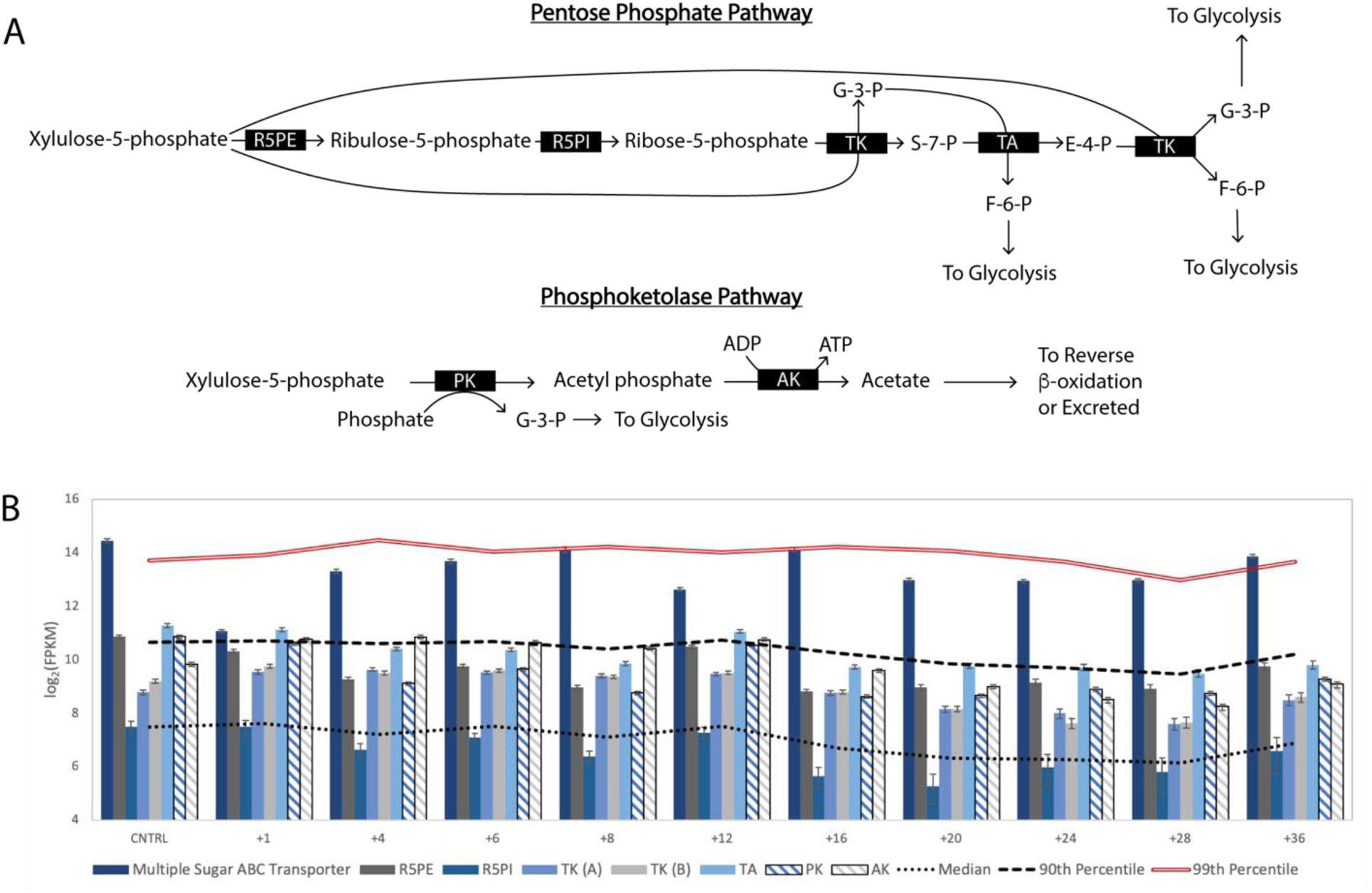
Predicted routes for xylose utilization by *Cand*. W. bifidus. (A) After xylose activation to xululose-5-phosphate, *Cand*. W. bifidus can consume pentoses via the pentose phosphate pathway or the phosphoketolase pathway. Key enzymes involved in xylose consumption include ribulose 5-phosphate epimerase (R5PE), ribose-5-phosphate isomerase (R5PI), transketolase (TK), transaldolase (TA), transketolase (TK), phosphoketolase (PK) and acetate kinase (AK). (B) Expression of genes involved with xylose utilization.

#### Predicted conservation of metabolic functions between *Cand*. W. bifidus and related organisms

We found that all of the UBA2727 genomes, *Shuttleworthia satelles*, and the LCO1.1 genome contained genes needed for the reverse β-oxidation pathway (**Supplementary Data File 2**). In the UBA2727, *S. satelles*, and *Cand*. W. bifidus genomes, the genes encoding ACAT, HAD, ACD, EtfA, and EtfB are clustered (**Fig. 3A-F**), whereas ECoAH is located on a different region of the genome. The presence of genes encoding each of these enzymes and their transcript abundance patterns in *Cand*. W. bifidus during MCFA production suggest that reverse β-oxidation and the electron-bifurcating ACD reaction are conserved and key features of chain elongation in each of these organisms. Genome comparisons revealed that the co-location of genes encoding ACD, EtfA, and EtfB is a common, although not essential, topology among known MCFA-producing organisms. Examples of other MCFA producers with this gene arrangement include *Megasphaera elsdenii* (**Fig. 3G**)*, Clostridium kluyveri* (**Fig. 3H**) and *Ruminococcaceae* bacterium CPB6 (**Fig.3I**).

While two routes of pentose utilization (the pentose phosphate and phosphoketolase pathways) are predicted to be present in the *Cand*. W. bifidus genome, none of the UBA2727 genomes, nor the *S. satelles* genome, contain a predicted phosphoketolase gene. This indicates that the use of multiple pentose consumption pathways may be a unique feature of *Cand*. W. bifidus when compared to other members of the UBA2727 cluster and to the nearest type strain. All of the UBA2727 genomes and *Cand*. W. bifidus contain genes for acetate production, including phosphotransacetylase and acetate kinase, indicating that all of the species in this genus may produce acetate as a product of anaerobic sugar metabolism.

The putative phosphoketolase from *Cand*. W. bifidus is similar to enzymes predicted to be present in other firmicutes and β-proteobacteria, including three species of *Megasphaera* (**Fig. S4**), a genus containing known MCFA producers. The phosphoketolase pathway (**Fig. 5A**), termed the “bifid shunt” (21) in a *Bifidobacterium* (22), provides an alternative to glycolysis and the pentose phosphate pathway for sugar utilization. In this pathway, the phosphoketolase enzyme splits xylulose-5-phosphate into glyceraldehyde-3-phosphate and acetyl-phosphate (23). Energy can then be conserved through the phosphorylation of ADP and production of acetate by acetate kinase (**Fig. 5A**). Overall, the phosphoketolase pathway can produce more ATP than the pentose phosphate pathway per mol of xylose (**Fig. S5**) and it directs carbon to acetate in addition to producing glycolysis intermediates (**Fig. 5A**).

#### Chain elongation by *Cand*. P. fermentans

We previously predicted that *Cand*. P. fermentans consumed lactate and produced MCFA when this microbiome was supplied with lignocellulosic biorefinery residues (8). In this study, we found that transcripts encoding enzymes predicted to function in the reverse β-oxidation pathway were among the most abundant in *Cand*. P. fermentans after the addition of lignocellulosic biorefinery residues (**Fig. 6A**). However, in the revised Cand*. P. fermentans* genome we find that the gene encoding the putative ECoAH protein is located near genes predicted to encode enzymes in the reverse β-oxidation cycle (**Fig. 3**), unlike the genomes of *Cand*. W. bifidus and related organisms. In addition, we find that the abundance of the transcript encoding the EcoAH protein correlates well with those of *Cand*. P. fermentans genes predicted to encode other enzymes in the reverse β-oxidation cycle (**Fig S6**). In *Cand*. P. fermentans, the genes encoding the predicted EtfA and EtfB proteins are not located next to those encoding the ACD enzyme. Instead, the *etfAB* genes are in a cluster with one that encodes a homologue of a putative prephenate dehydrogenase (PRDH), an enzyme that catalyzes an oxidative decarboxylation in the shikimate pathway for tyrosine biosynthesis (24) (**Fig. 3**). A pairwise gene expression analysis of transcript levels encoding the ACD, PRDH, EtfA, and EtfB proteins (**Fig. 7**) showed that ACD had strong correlations with EtfA (r^2^=0.93) and EtfB (r^2^=0.91), suggesting co-regulation of ACD and the electron transport flavoproteins, and supporting a role for an electron-bifurcating ACD in the reverse β-oxidation cycle of *Cand*. P. fermentans, as we also predict for *Cand*. W. bifidus.

**Figure 6.**
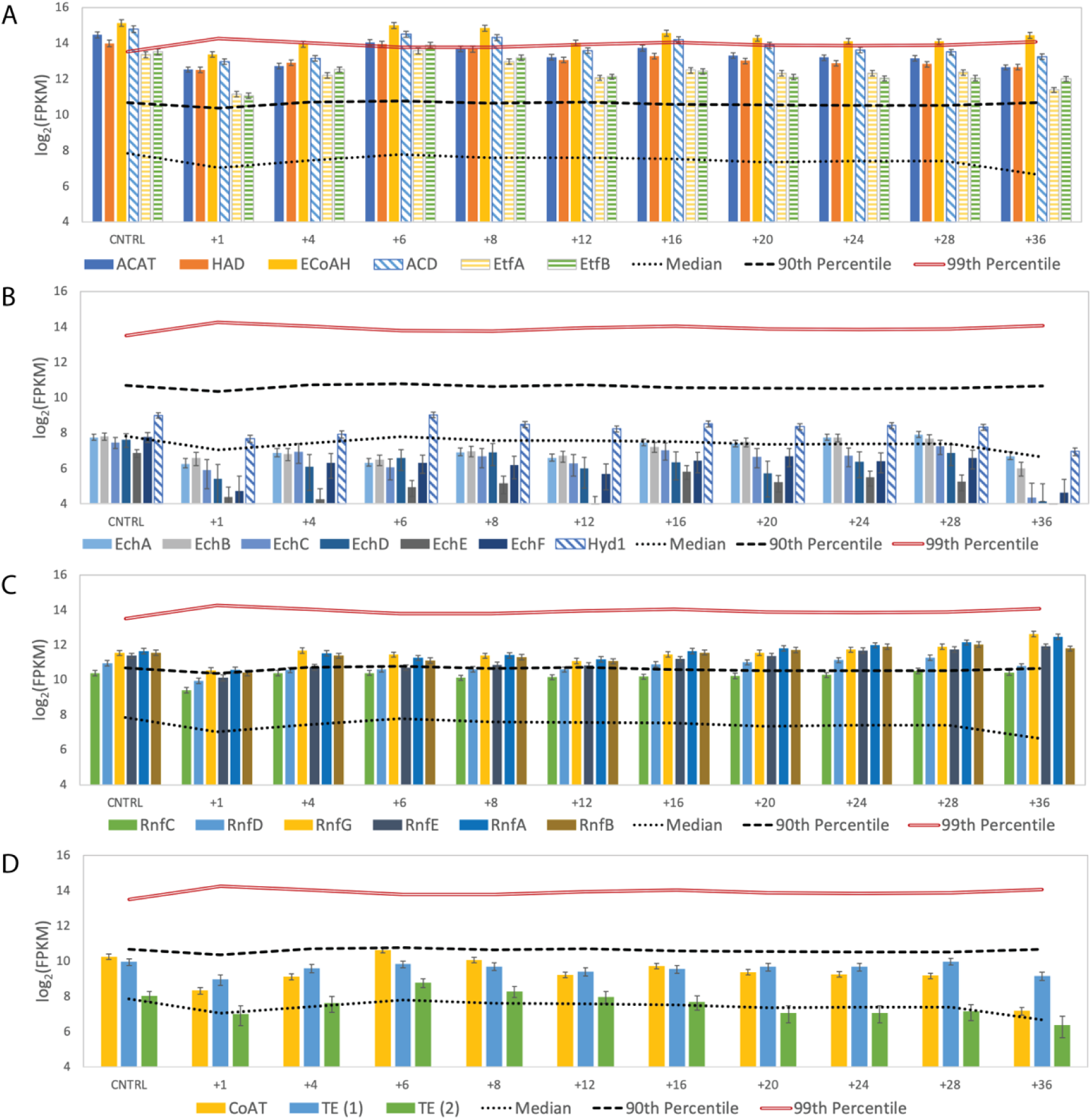
Expression of chain elongation genes by *Cand*. P. fermentans. Genes expression data is shown for genes encoding enzymes used for (A) reverse β-oxidation, (B) the RNF complex, (C) hydrogenases, (D) and terminal enzymes of reverse β-oxidation. The roles of enzymes in chain-elongation and enzyme definitions are shown in Fig 2.

**Figure 7.**
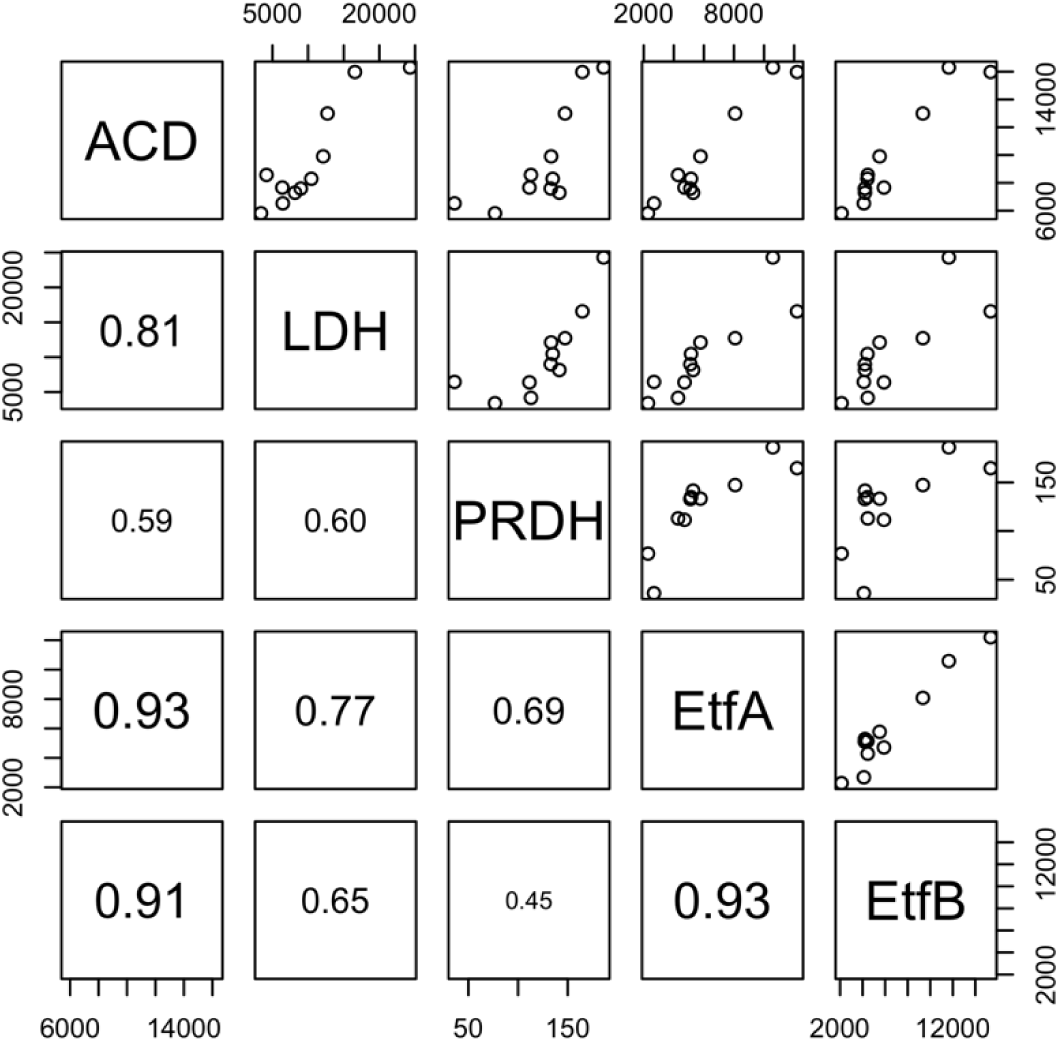
Pairwise linear correlation analyses for expression of *Cand* P. fermentans genes encoding lactate dehydrogenase (LDH), acetyl-CoA dehydrogenase (ACD), prephenate dehydrogenase (PRDH), Electron transfer flavoprotein A (EtfA), and electron transfer flavoprotein B (EtfB). FPKM values are indicated on x- and y-axes. Coefficients of determination are shown in bottom-left panels.

The revised genome sequence of *Cand*. P. fermentans also contained complete sets of genes for subunits of the RNF and the Ech complexes, a characteristic also shared with *Cand*. W. bifidus. The transcript abundance of *Cand*. P. fermentans genes encoding subunits of the RNF complex were at or near the 90^th^ percentile throughout the time series analyzed in this experiment (**Fig. 6B**), whereas the transcript abundance for the genes predicted to encode the *Cand*. P. fermentans Ech hydrogenase were below the median at most time points after the addition of lignocellulosic biorefinery residues (**Fig. 6C**). Furthermore, the other hydrogenase predicted to be present in *Cand*. P. fermentans, Hyd1, had transcript abundances that were higher than those of genes encoding the Ech hydrogenase complex. These observations suggest that energy conservation in *Cand. P. fermentans* primarily occurs via generation of an ion motive force by the RNF complex. However, hydrogen production via Hyd1 could be important to maintain redox balance during chain elongation by *Cand*. P. fermentans.

When considering hydrolysis of the elongated acyl-CoA molecule, the *Cand*. P. fermentans genome encodes a CoAT and two thioesterases, each of which were expressed at or above median levels when compared to other transcripts analyzed during this experiment (**Fig. 6D**). The abundance of the transcripts encoding CoAT and one of the thioesterases (TE.2; **Fig. S6**) have a higher correlation with those of predicted enzymes in the reverse β-oxidation pathway than the transcripts of the other thioesterase (TE.1; **Fig. S6**). As it was the case with TE.1 in *Cand*. W. bifidus, an amino acid sequence analysis of the TE.2 from *Cand*. P. fermentans showed that it belongs to the hotdog-fold superfamily of thioesterases. Specifically, the TE.2 *Cand* P. fermentans thioesterase is predicted to be a PaaI family enzyme, which includes the medium-chain acyl-CoA thioesterase II of *Escherichia coli* (25). Thus, the transcriptomic analysis suggests that CoAT and TE.2 may both participate in acyl chain release during MCFA synthesis in *Cand*. P. fermentans.

#### Cand. *P. fermentans* is predicted to use multiple routes for glycerol metabolism

Glycerol is known to be a significant carbon source in the lignocellulosic biorefinery residues used in this and earlier studies (8). Analysis of the reactor media showed that all the glycerol was removed within the first 6 hours (**Fig. S2**) suggesting it is a favored carbon source for one or more of the microbes in this microbiome. The most highly expressed gene by *Cand*. P. fermentans at several time points after addition of lignocellulosic biorefinery residues encodes a predicted glycerol transporter (**Fig. 8B**), suggesting that glycerol is rapidly transported by this organism.

**Figure 8.**
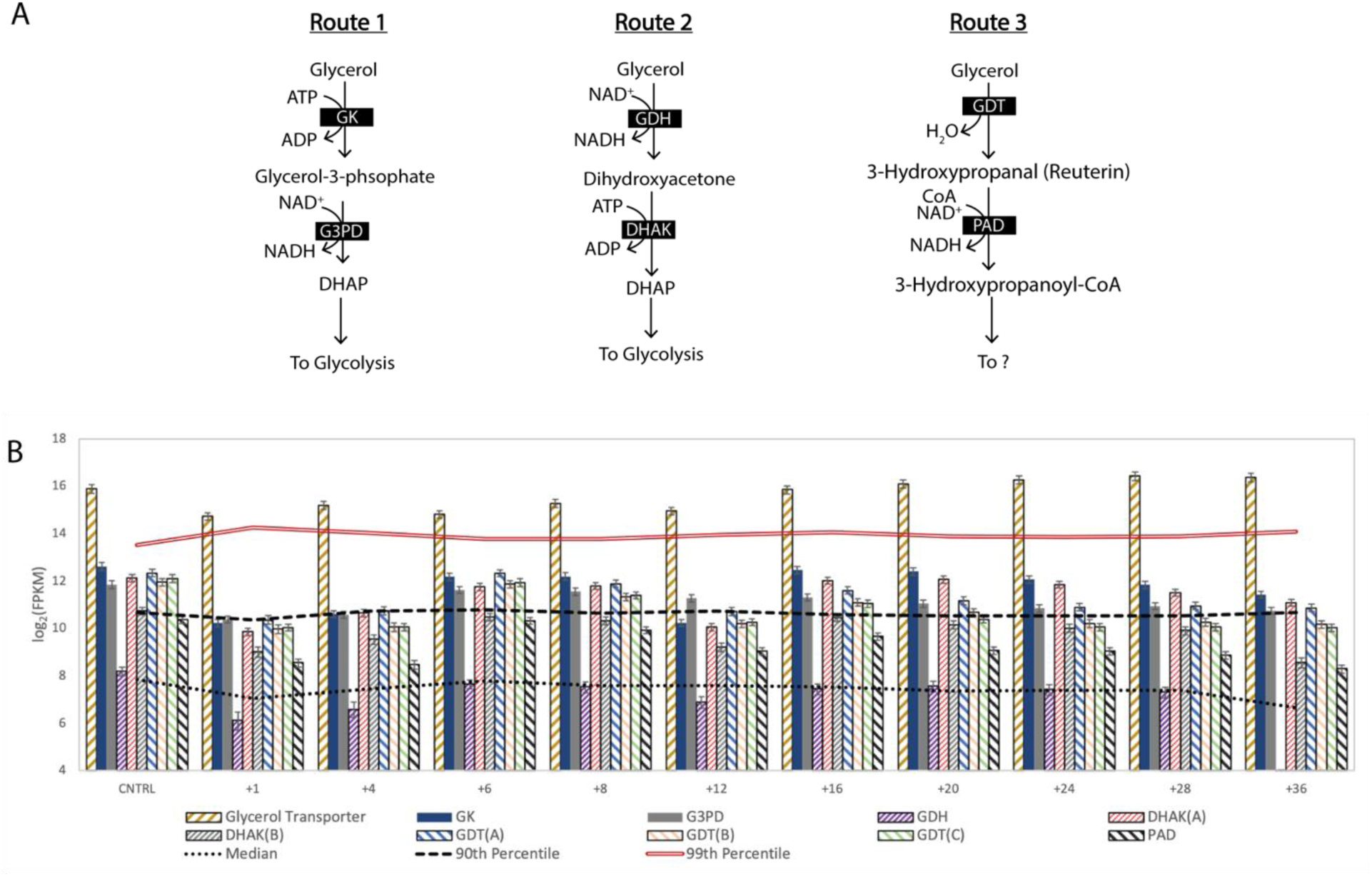
Glycerol utilization by *Cand*. P. fermentans. (A) *Cand*. P. fermentans contains genes for three glycerol utilization routes. Key enzymes in these pathways include glycerol kinase (GK), glycerol-3-phosphate dehydrogenase (G3PD), glycerol dehydrogenase (GDH), dihydroxyacetone kinase (DHAK), glycerol dehydratase (GDT), and propionaldehyde dehydrogenase (PAD). (B) Expression of genes for glycerol transport and utilization. Subunits of a single enzyme complex are indicated in parentheses.

To assess how *Cand*. P. fermentans metabolizes glycerol, we monitored transcript abundance of genes predicted to be involved in this process. The revised *Cand*. P. fermentans genome also predicts that this organism contains enzymes to metabolize glycerol, and the time series transcriptomics data showed that it expressed genes for three putative glycerol conversion pathways (**Fig. 8**). The first route predicted to be active in *Cand*. P. fermentans (**Fig. 8A**, Route 1) uses an ATP-dependent glycerol kinase, in which the resultant glycerol-phosphate is oxidized to produce dihydroxy-acetone phosphate (DHAP). The second predicted route in *Cand*. P. fermentans (**Fig. 8A**, Route 2) uses glycerol dehydrogenase to oxidize glycerol to dihidroxyacetone before it is phosphorylated to DHAP. In a third predicted route in *Cand*. P. fermentans (**Fig. 8A**, Route 3), glycerol is converted to 3-hydroxypropanal (reuterin), a compound proposed to inhibit growth of other microbes by inducing oxidative stress (26). All of the above predicted glycerol utilization genes, with the exception of glycerol dehydrogenase, are expressed above median levels throughout the 36-hour time series (**Fig. 8B**). Thus, it is possible that multiple or even all three pathways (**Fig. 8A**) play a role in glycerol metabolism in *Cand*. P. fermentans.

In addition to expressing genes encoding pathways for multiple glycerol utilization routes, *Cand*. P. fermentans contains a gene cluster encoding predicted propanediol utilization body proteins along with genes encoding the multiple subunits of a glycerol dehydratase (**Fig. 9**). Thus, it is possible that these predicted protein microcompartments may help protect *Cand*. P. fermentans from a toxin like reuterin, or others, that are potential products of glycerol metabolism by one or more routes in this organism.

**Figure 9.**
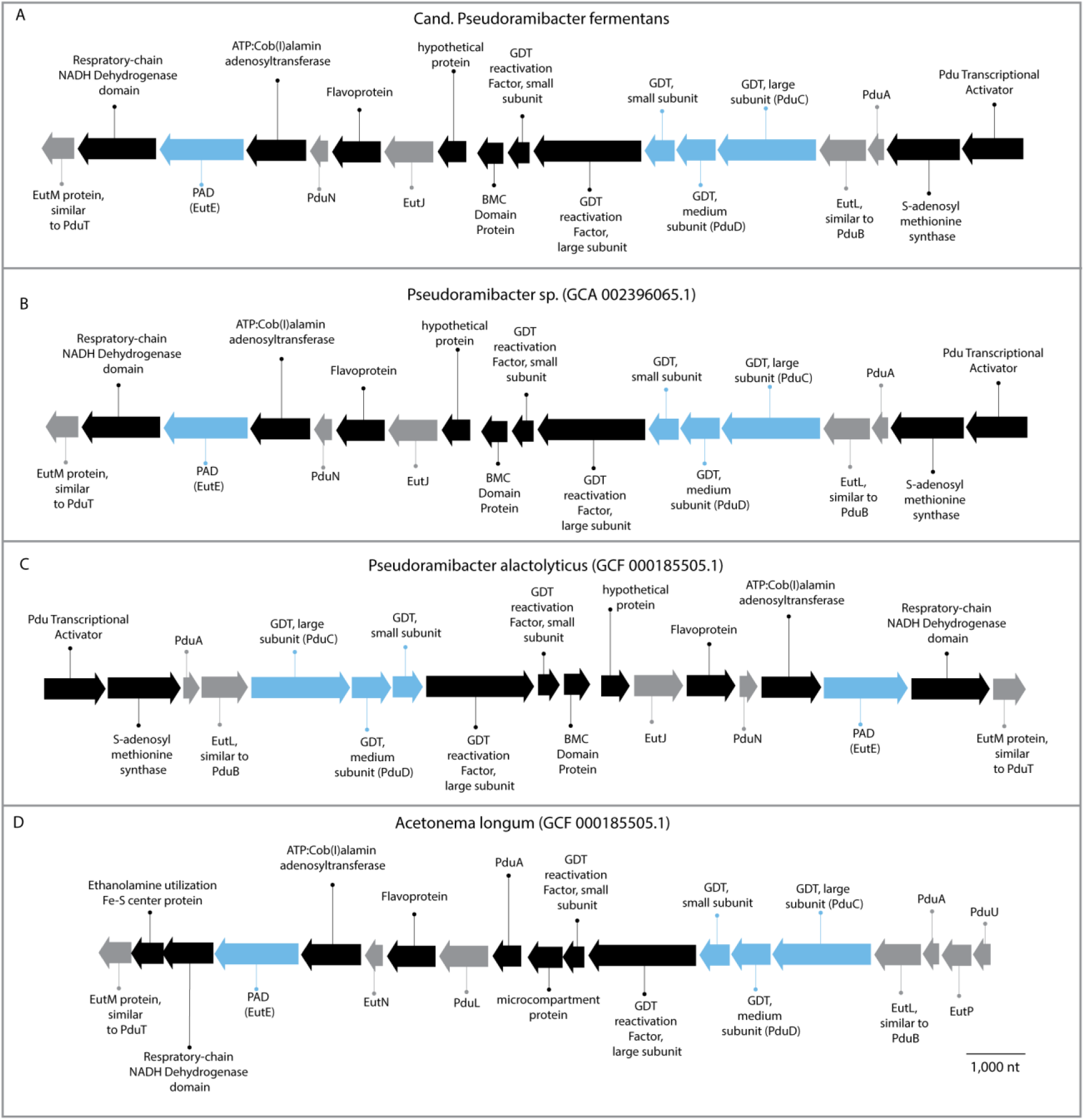
Glycerol utilization gene clusters in *Cand*. P. fermentans (A), related *Pseudoramibacter* species (B-C), and *Acetonema longum* (D). Blue shading indicates a gene predicted to encode an enzyme involved in Route 3 (Fig 6A) and grey shading indicates a gene predicted to encode a microcompartment protein.

Overall, our data provides new evidence that *Cand*. P. fermentans directs glycerol to central carbon metabolism and potentially produces toxic compounds that may impact the growth and abundance of other members in this microbiome. One possible outcome of glycerol metabolism to intermediates that can be further transformed via glycolysis (**Fig. 8**) is its conversion to MCFA. A free energy analysis of glycerol conversion to butyrate, hexanoate, and octanoate shows that these are all energetically feasible reactions when hydrogen is produced to maintain internal redox balance (**Table 2**).

**Table 2.**
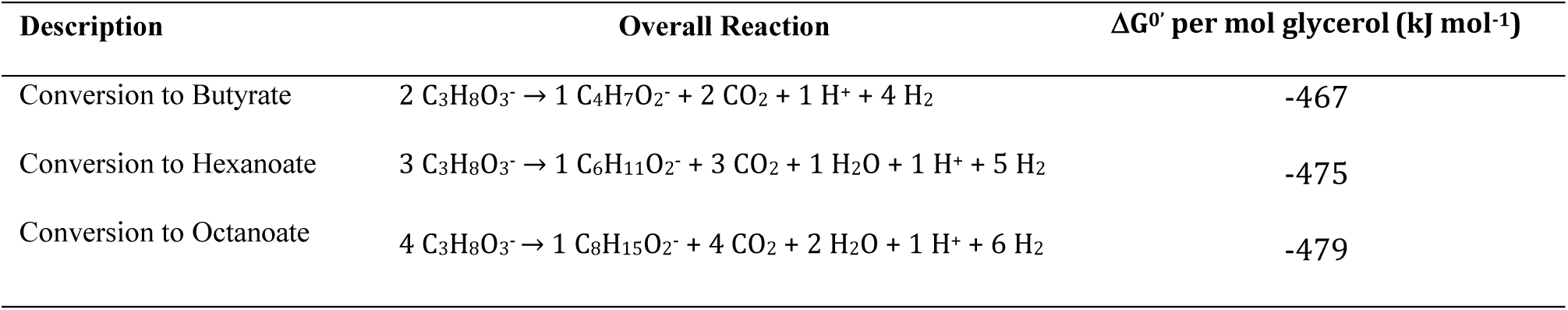
Thermodynamics of glycerol conversion to butyrate, hexanoate, and octanoate at high and low hydrogen partial pressures

| Description | Overall Reaction | $\Delta G^{\circ}$ per mol glycerol (kJ mol<sup>-1</sup>) |
| --- | --- | --- |
| Conversion to Butyrate | $2 \text{ C}_3\text{H}_8\text{O}_3^- \rightarrow 1 \text{ C}_4\text{H}_7\text{O}_2^- + 2 \text{ CO}_2 + 1 \text{ H}^+ + 4 \text{ H}_2$ | -467 |
| Conversion to Hexanoate | $3 \text{ C}_3\text{H}_8\text{O}_3^- \rightarrow 1 \text{ C}_6\text{H}_{11}\text{O}_2^- + 3 \text{ CO}_2 + 1 \text{ H}_2\text{O} + 1 \text{ H}^+ + 5 \text{ H}_2$ | -475 |
| Conversion to Octanoate | $4 \text{ C}_3\text{H}_8\text{O}_3^- \rightarrow 1 \text{ C}_8\text{H}_{15}\text{O}_2^- + 4 \text{ CO}_2 + 2 \text{ H}_2\text{O} + 1 \text{ H}^+ + 6 \text{ H}_2$ | -479 |

#### *Cand*. P. fermentans is predicted to use an electron-confurcating lactate dehydrogenase for MCFA production

We previously proposed that one member of this MCFA-producing microbiome consumes lactate produced by other community members (8). The *Cand*. P. fermentans genome assembled in this study contains a gene cluster encoding a lactate transporter (lactate permease), lactate dehydrogenase (LDH), and ACD (**Fig. 3J**). All three of these genes are expressed above the 90^th^ percentile at all time points after the addition of lignocellulosic biorefinery residue (**Fig. 10**), supporting their role in MCFA production from lactate.

**Figure 10.**
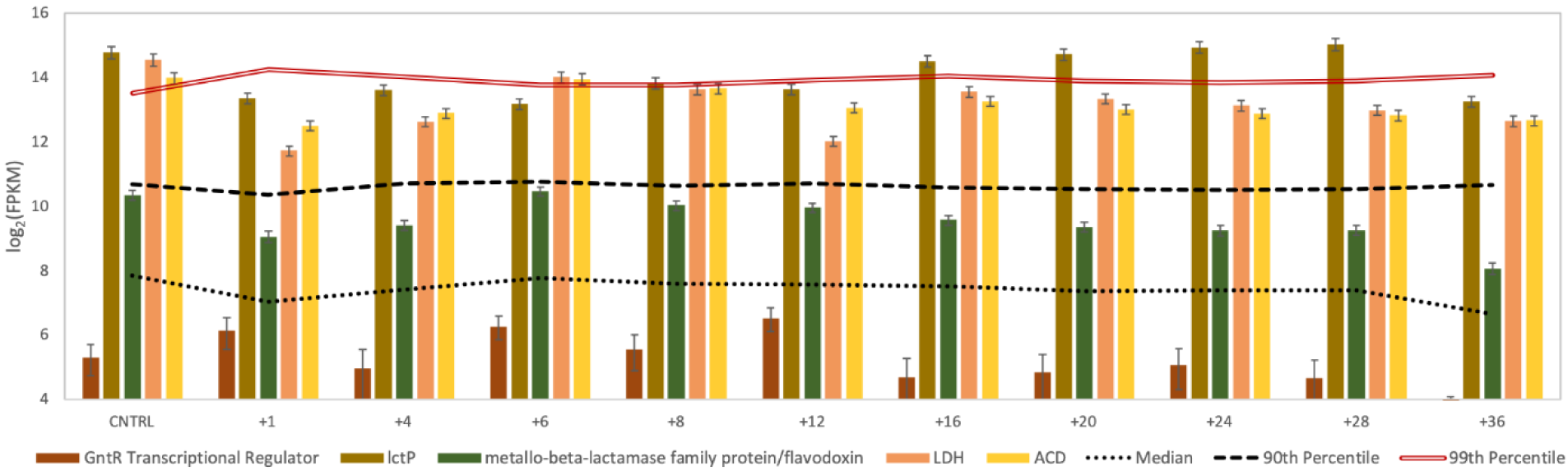
Expression of genes in a lactate utilization gene cluster in *Cand* P. fermentans. Genes include a transcriptional regulator, lactate permease (*lctP*), a potential flavodoxin, lactate dehydrogenase (LDH), and acetyl-CoA dehydrogenase (ACD).

The amino acid sequence of the predicted *Cand*. P. fermentans LDH in this gene cluster is most closely related to FAD-binding oxidoreductases contained in genomes from several related organisms, including other *Firmicutes* and organisms in the *Fusobacterium* phylum (**Supplementary Data File 8**). This *Cand*. P. fermentans LDH clusters with proteins from organisms that consume lactate anaerobically (**Fig. 11**), including two bacteria known to convert lactate to MCFA, *Ruminococcaeae* CPB6 and *Megasphaera elsdenii* (5, 7), and with the LDH from *A. woodii* that uses electron confurcation to couple lactate and ferredoxin oxidation with NAD^+^ reduction to overcome the thermodynamic bottleneck of lactate oxidation (27). Pairwise correlation analyses of transcript abundance (**Fig. 8**) for this *Cand*. P. fermentans LDH with EtfA (r^2^=0.77) and EtfB (r^2^=0.65) suggest a role for this dehydrogenase and the electron transfer flavoproteins in anaerobic lactate oxidation. Therefore, we hypothesize that in *Cand*. P. fermentans, the electron transfer flavoproteins EtfA and EtfB have a role in both electron bifurcation by ACD in the reverse β-oxidation cycle and in electron confurcation with LDH.

#### Predicted conservation of metabolic functions between *Cand. P*. fermentans and related organisms

The organization of genes encoding for the complete reverse β-oxidation pathway in *Cand*. P. fermentans is similar to that found in the other two genomes available for *Pseudoramibacter* organisms (**Fig. 3**). In all three cases, the gene encoding ECoAH clusters with other genes in the reverse β-oxidation pathway, while the *etfAB* genes are found in a separate region of the genome. In all three organisms, the reverse β-oxidation genes are located near the genes encoding fatty acid biosynthesis (**Fig. 3**). While others have suggested a potential role for fatty acid biosynthesis genes in MCFA production (28), our data do not provide support for this hypothesis since transcripts for gene encoding enzymes involved in reverse β-oxidation were orders of magnitude more abundant than fatty acid biosynthesis genes (**Fig. S7**).

**Figure 11.**
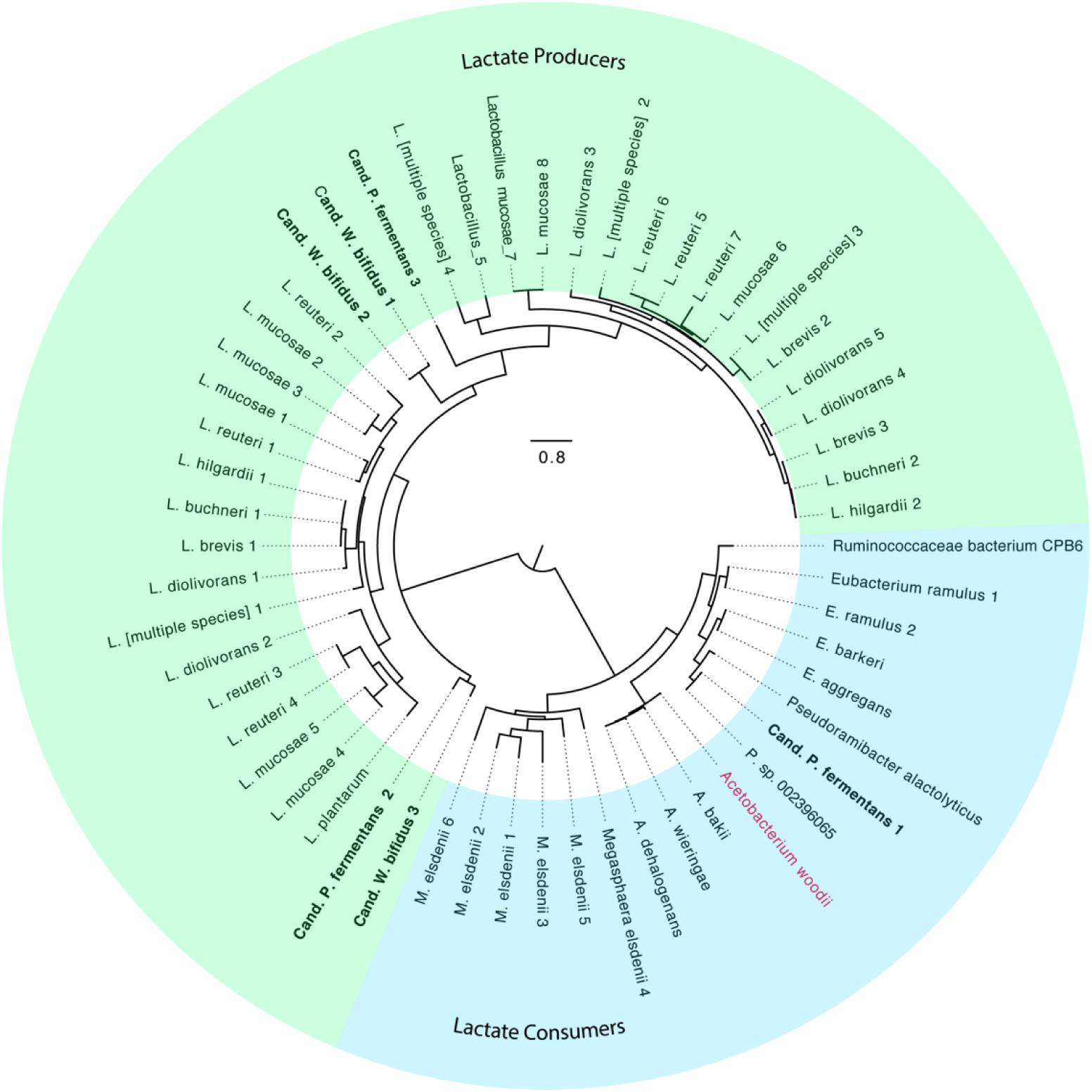
A maximum-likelihood phylogenetic tree of lactate dehydrogenase amino acid sequences among different members of the *Firmicutes* phylum. Anaerobic lactate consumers contain lactate dehydrogenase that form a distinct cluster from lactate producers, such as those found among the *Lactobacilli*.

The ability of Cand. *P. fermentans* to metabolize glycerol appears to be common in the *Pseudoramibacter* genus, as genes for the three proposed routes of glycerol metabolism in *Cand*. P. fermentans are also found in the other two *Pseudoramibacter* genomes. This is also the case for the presence of genes predicted to produce protein microcompartments in a cluster with genes encoding glycerol dehydratase (**Fig. 9**). The production of polyhedral protein microcompartments thought to encase a diol dehydratase when consuming 1,2-propanediol has been described in other organisms, such as *Salmonella enterica* (a *γ-Proteobacterium*) and *Acetonema longum* (**Fig. 9D**), a member of the *Negativicutes* class within the *Firmicutes* phylum (29, 30). In addition to the two other *Pseudoramibacter* genomes which contain identical glycerol gene clusters to *Cand*. P. fermentans (**Fig. 9B-C**), a similar glycerol utilization cluster was identified in *A. longum* (**Fig. 9D**).

### Summary descriptions of novel organisms

#### Description of genus *Candidatus* Weimerbacter, genus, nov.

(Named for Paul J. Weimer, a pioneer in using rumen microbes to produce valuable chemicals). The genus *Weimerbacter* belongs to the *Lachnospiraceae* family. This genus contains bacteria from the cattle rumen and a lignocellulose-fed anaerobic bioreactor. An isolate from this genus was obtained as part of the Hungate 1000 project (10). Species within this genus contain genes needed for utilization of sugars, acetate production, lactate production, reverse β-oxidation, pyruvate flavodoxin oxidoreductase, the RNF complex, and hydrogen production. Our analysis predicts that organisms in the *Weimerbacter* genus can produce MCFA when using xylose as an organic substrate. A predicted *Weimerbacter* energy conserving mechanism used during MCFA production involves an electron-bifurcating acyl-CoA dehydrogenase (ACD); the reduced ferredoxin derived from this activity is used by either the RNF complex or the Ech hydrogenase complex for ion translocation that contributes to the creation of an ion motive force for ATP production.

#### Description of *Candidatus* Weimerbacter bifidus, sp. nov. (uses the bifid shunt, bifidus)

*Candidatus* Weimerbacter bifidus is represented by the LCO1.1 metagenome assembled genome (SAMN12235634). When a bioreactor community is fed lignocellulosic biorefinery residues, *Cand*. W. bifidus is predicted to use the phosphoketolase pathway (bifid shunt) to degrade xylose and produce butyrate and medium-chain fatty acids via reverse β-oxidation. *Cand*. W. bifidus is predicted to maintain redox balance by using a ferredoxin-dependent hydrogenase or an energy conserving Ech hydrogenase to produce hydrogen. While other organisms within the *Cand*. Weimerbacter genus contain genes for pentose degradation via the oxidative pentose phosphate pathway, *Cand*. W. bifidus is unique in containing genes for the bifid shunt.

#### Description of *Candidatus* Pseudoramibacter fermentans, sp. nov. (fermentans, found in fermentation bioreactor)

*Candidatus* Pseudoramibacter fermentans is represented by the metagenome assembled genomes EUB1.1 (SAMN12235633) and *Pseudoramibacter* sp. (GCA 002396065.1), and it is differentiated from the *P. alactolyticus* species by an average nucleotide identity lower than 95%. Unlike *P. alactolyticus*, both the EUB1.1 and the *Pseudoramibacter* sp. (GCA 002396065.1) contain genes that code for a proton-translocating energy conserving hydrogenase (EchABCDEF). When lignocellulosic biorefinery residues are fed to a bioreactor community, *Cand*. P. fermentans is predicted to consume glycerol that is present in the residue and lactate that can be produced by other microbial community members. This species is predicted to produce MCFA, via reverse β-oxidation, using lactate and glycerol as the organic substrates. *Cand*. P. fermentans contains genes for multiple glycerol utilization routes, including conversion of glycerol to reuterin. Lactate utilization is predicted to involve an electron-confurcating lactate dehydrogenase (LDH). In addition, an electron-bifurcating acyl-CoA dehydrogenase (ACD) is predicted to be used for energy conservation, with reduced ferredoxin produced by this enzyme used by the RNF complex for the creation of an ion motive force to support ATP production. *Cand*. P. fermentans is predicted to produce hydrogen via a ferredoxin-dependent hydrogenase as a mechanism for balancing internal redox conditions.

### Concluding remarks

Our results reveal several previously unexplored metabolic and energetic features of chain-elongating bacteria. Multi-omic analysis of *Cand*. W. bifidus suggest that this organism may have a previously undescribed ability to use both the pentose phosphate and the phosphoketolase pathways for pentose consumption and production of acetyl CoA that is needed for MCFA synthesis. Further, both *Cand*. W. bifidus and *Cand*. P. fermentans may use multiple hydrogenases, including a proton-translocating energy conserving hydrogenase (EchABCDEF) to support MCFA production. Although both chain elongators contained genes for the RNF complex, the transcriptomic evidence suggests that this complex may be more important for generating an ion motive force in *Cand*. W. bifidus. Our data also predict that *Cand*. P. fermentans uses several routes to consume glycerol as a carbon source, with potentially toxic intermediates sequestered in protein microcompartments, and a thermodynamic analysis supports the ability of *Cand*. P. fermentans to produce MCFA from this substrate. Finally, our data implicates an electron-confurcating LDH in providing carbon skeletons needed to support MCFA production from lactate by *Cand*. P. fermentans. Further work is necessary to elucidate the implications of the genomic features uncovered in this study on the bioenergetics of MCFA production, to assess whether 19 similar processes are involved in the production of MCFA from other substrates, and to develop a better understanding of microbial pathways for production of additional valuable products from renewable organic materials.

## METHODS

### Bioreactor operation

We operated a bioreactor containing 150 mL of liquid. Lignocellulosic biorefinery residues, prepared as described previously (2) was added into the bioreactor and reactor liquid was pumped out of the bioreactor every hour to maintain a residence time in the reactor of 6 days. The pH of the reactor was controlled at 5.5 by adding 5M KOH through a pump attached to a pH controller. The temperature of the bioreactor was maintained at 35 °C using a water bath. For the 36-hour time series experiment, 28 mL of liquid was removed from the reactor and 28 mL of lignocellulosic biorefinery residues was added one time, bringing the starting liquid volume to 150 mL.

### Reactor sampling

Prior to feeding the reactor with 28 mL of lignocellulosic biorefinery residues, samples were collected for metagenomic and metatranscriptomic analyses. Samples for DNA sequence analysis were collected in 2 mL centrifuge tubes and centrifuged at 10,000 g for 10 minutes. After decanting the supernatant, cell pellets were stored at −80 °C until DNA was extracted. Samples for RNA were collected in 2 mL centrifuge tubes and centrifuged at 10,000 g for 1.5 minutes. After decanting the supernatant, samples for RNA were flash frozen in liquid nitrogen then stored at −80 °C until RNA was extracted. Three samples for RNA extraction and sequencing were collected as a control prior to adding conversion residue. At each time point after adding lignocellulosic biorefinery residues, one sample was collected for RNA extraction and the supernatant from these samples was used for HPLC and GC-MS analyses, as described previously (2).

### DNA sequencing

DNA was extracted from biomass samples using a phenol-chloroform extraction method described previously (8). For all metagenomic samples (Days 96, 120, 168, 252, and 378), sequencing was performed with an Illumnia HiSeq 2500 sequencer to generate 2 x 250 bp reads. For the Day 378 sample, PacBio sequencing and library preparation was performed by the Joint Genome Institute. The sequence library was prepared using a PacBio 10 kb low input library preparation and libraries were sequenced with a PacBio Sequel. The raw DNA sequences are available at the National Center for Biotechnology Information Sequencing Read Archive database under BioProject PRJNA393345 for Days 96 and 120, and BioProject PRJNA535528 for Days 168, 252, and 378.

### Metagenomic Analysis

Quality checking and trimming of Illumina DNA reads was performed using sickle (31). The resulting reads from Days 96, 120, 168, 252, and 378 were co-assembled using metaspades version 3.12.00 (32). The assembled contigs were binned using Anvio version 5 (33). After binning, draft genomes were gap-filled with the PacBio reads from Day 378 using PBSuite version 15.8.24 (34). Quality checking was performed on the gap-filled draft genomes using CheckM (35). Draft genomes were annotated with Metapathways version 2.5 (36). Taxonomic assignments based on single-copy marker genes were made with GTDB tool kit (9) and phylogenetic trees were constructed using RAxML (37). Average nucleotide identities were calculated with JSpecies using the ANIb algorithm (38).

### Metatranscriptomic analysis

RNA was extracted and cDNA was synthesized and sequenced as described previously (8). Quality checking and trimming of raw reads was performed with sickle (39), and rRNA reads were removed with SortMeRNA (40). Reads from each time point were mapped to open reading frames in the MAGs with CuffLinks (41). The raw RNA sequences are available at the National Center for Biotechnology Information Sequencing Read Archive database under BioProject PRJNA535528.

## ACKNOWLEDGMENTS

This work was funded by the DOE Great Lakes Bioenergy Research Center (DOE BER Office of Science DE-SC0018409). M.J.S. is supported by the National Science Foundation Graduate Research Fellowship Program under grant No. DGE-1256259. The authors thank the University of Wisconsin Biotechnology Center and the U.S. Department of Energy Joint Genome Institute for library preparation and sequencing. The work conducted by the U.S. Department of Energy Joint Genome Institute, a DOE Office of Science User Facility, is supported by the Office of Science of the U.S. Department of Energy under Contract No. DE-AC02-05CH11231. The authors also thank Pamela Camejo and Francisco Moya for assistance with RNA sequencing and data analyses.

## SUPPLEMENTARY FIGURES

**Figure S1.**
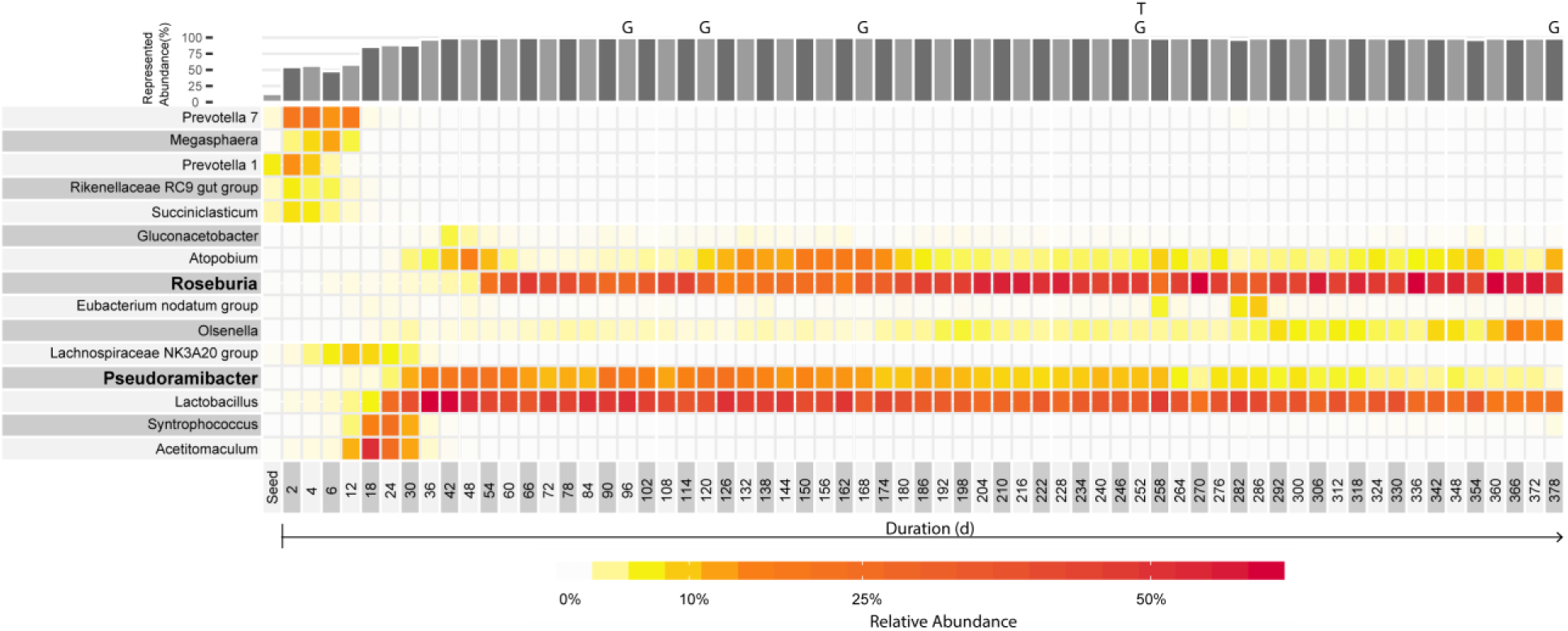
Relative abundance of bacteria in the bioreactor based on 16S rRNA gene amplicon sequencing. With the short 16S rRNA gene fragments, *Cand*. W. bifidus was classified as a member of the *Roseburia* genus, whereas *Cand*. P. fermentans was correctly classified as a member of the *Pseudoramibacter* genus. The first column shows results from the acid digester sludge (“seed”) used for reactor inoculum. The duration after starting the bioreactor is shown on the x-axis and genera names are provided on the y-axis. The bar plot above the heatmap shows the sum of abundance represented in the heatmap. Colors in the heatmap indicate relative abundance with higher abundance indicated by red color intensity. Samples corresponding to metagenomic and metatranscriptomic samples analyzed in this study are shown with “G” indicating a metagenomic sample and “T” indicating the time point used for the time-series metatranscriptomic analysis. The 16S-based abundance for the first 252 days was previously published in Scarborough et al. 2018 (2).

**Figure S2.**
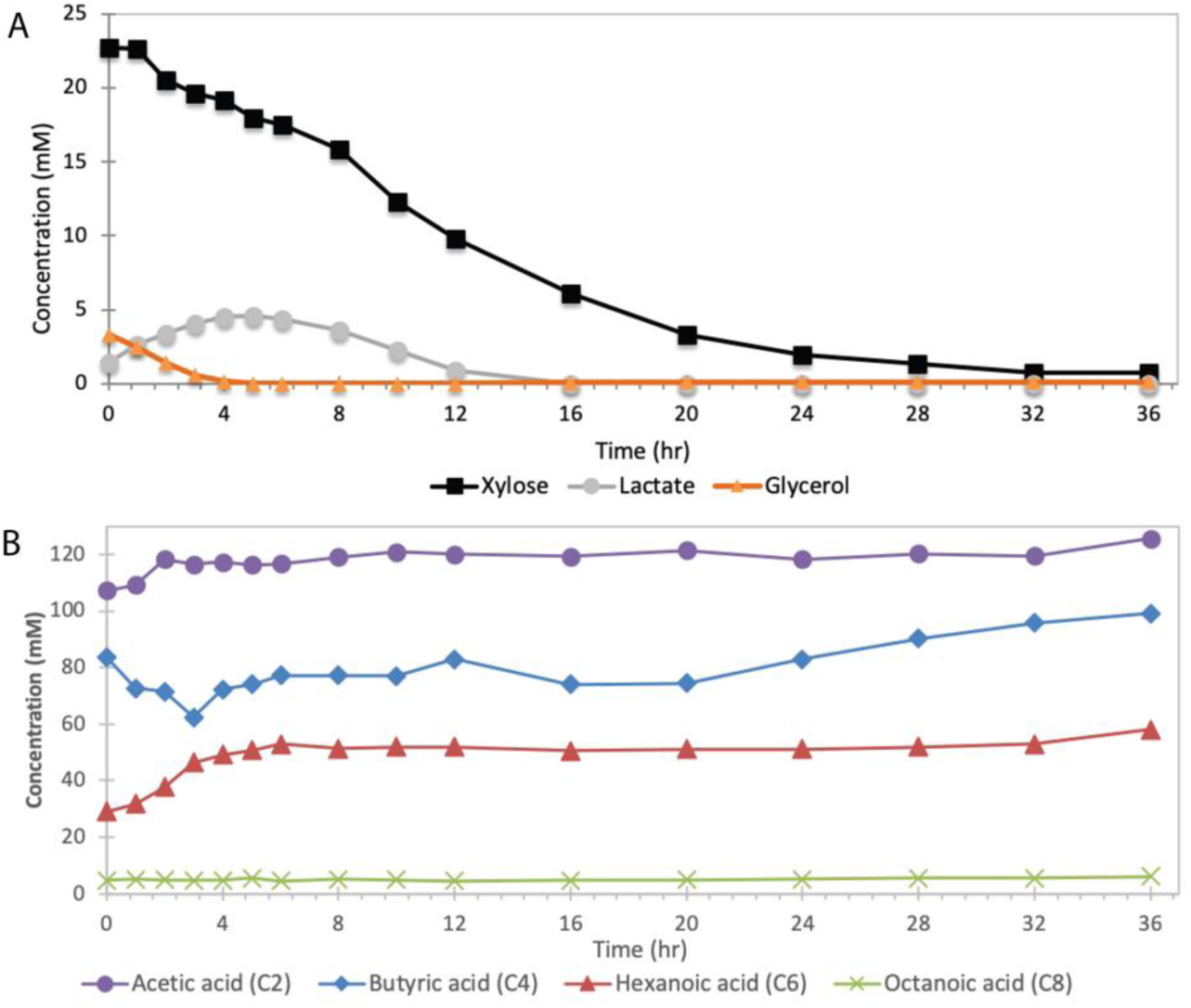
Extracellular metabolites for 36 hours after addition of lignocellulosic biorefinery residues. (A) Xylose, lactate, and glycerol are all compounds that are consumed during the course of the experiment. (B) The reactor produced various carboxylic acid end products, including acetic acid, butyric acid, hexanoic acid, and octanoic acid.

**Figure S3.**
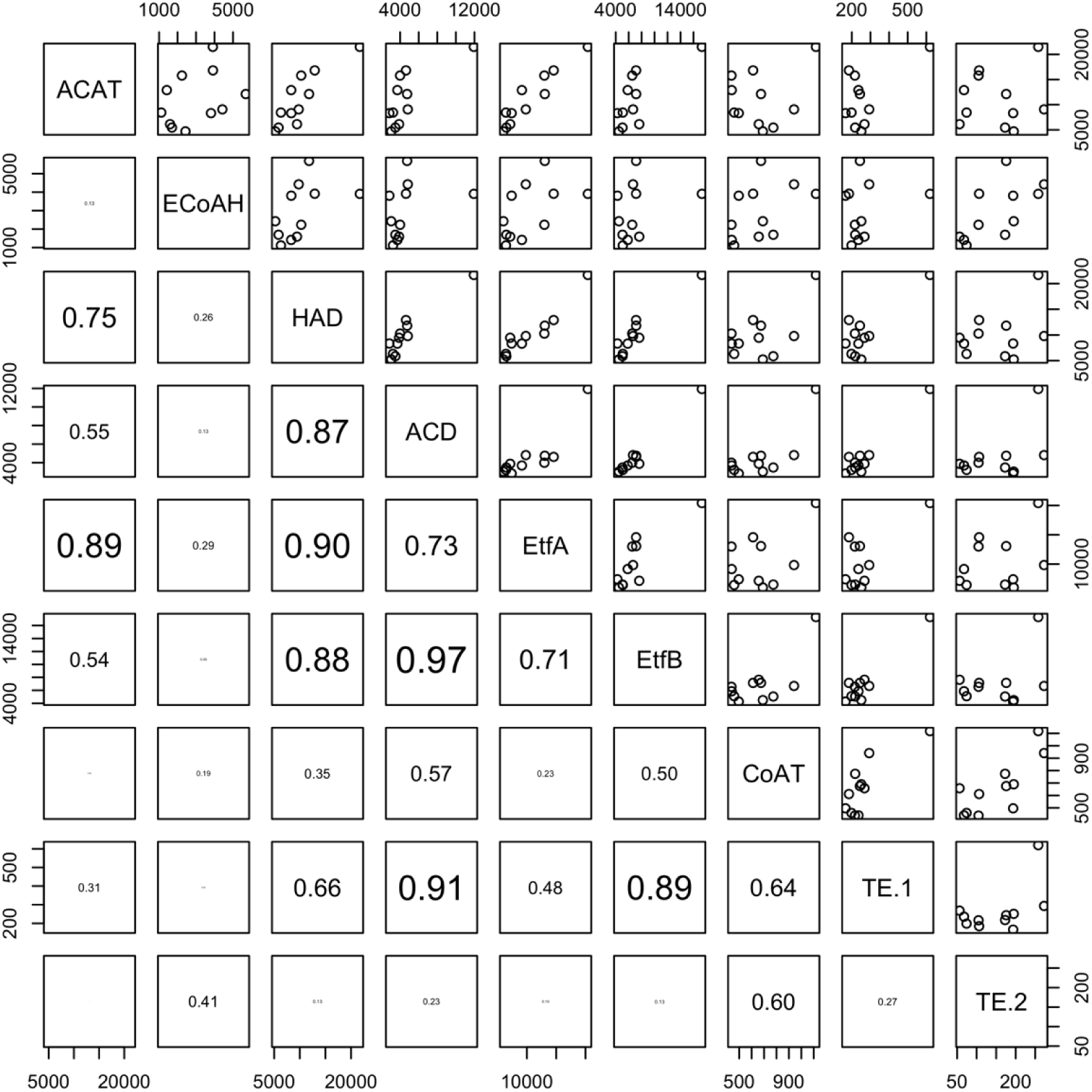
Pairwise comparison of transcript abundance for reverse β-oxidation genes in *Cand*. W. bifidus. Coefficients of determination are shown in bottom-left panels.

**Figure S4.**
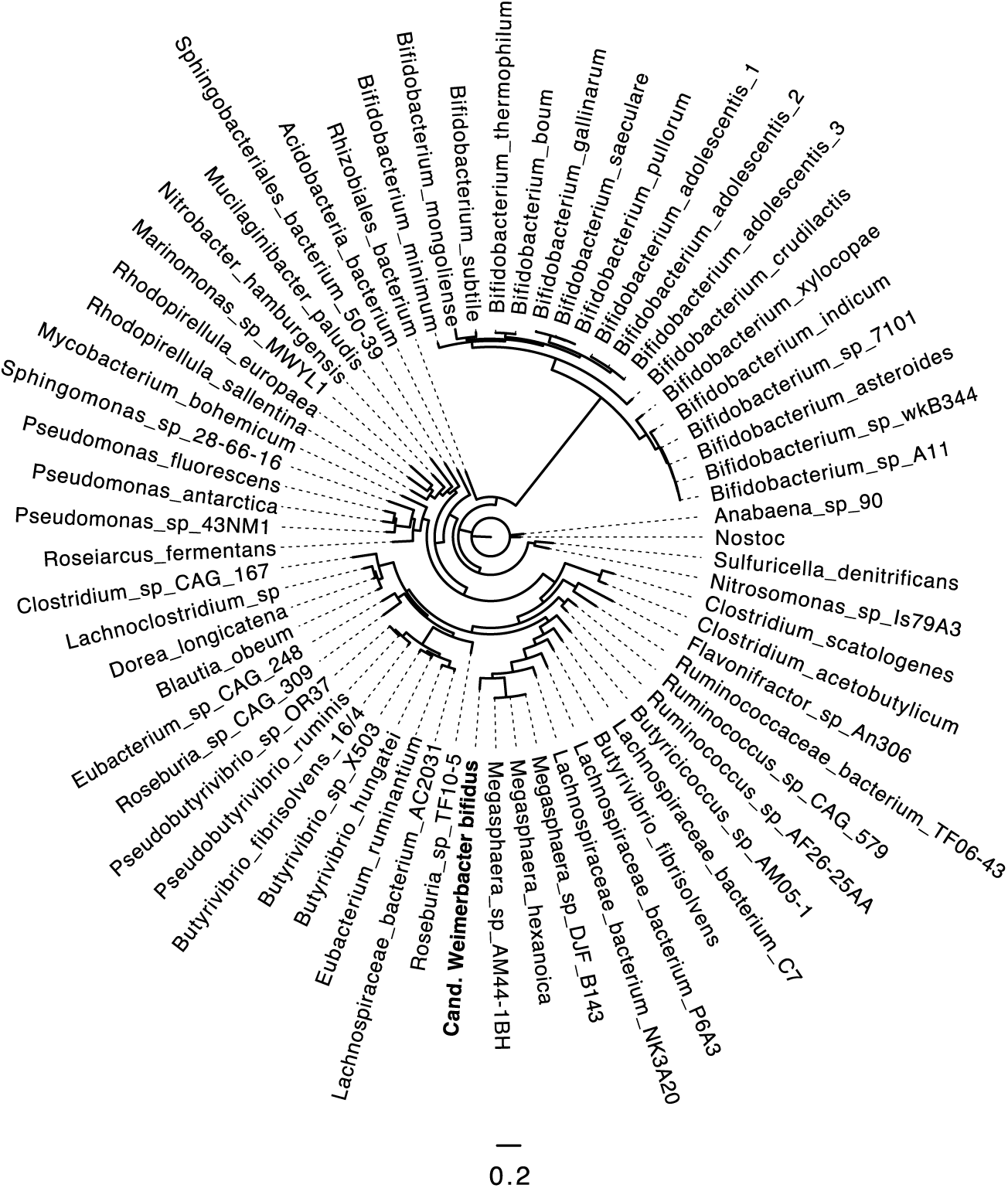
A maximum-likelihood phylogenetic tree of the phosphoketolase enzyme across several phyla of bacteria. Sequences across several phyla were selected and show a distinct cluster of the amino acid sequences for *Bifidobacterium* species. The sequence from *Cand*. W. bifidus clustered with those from members of the chain-elongating genus *Megasphaera*.

**Figure S5.**
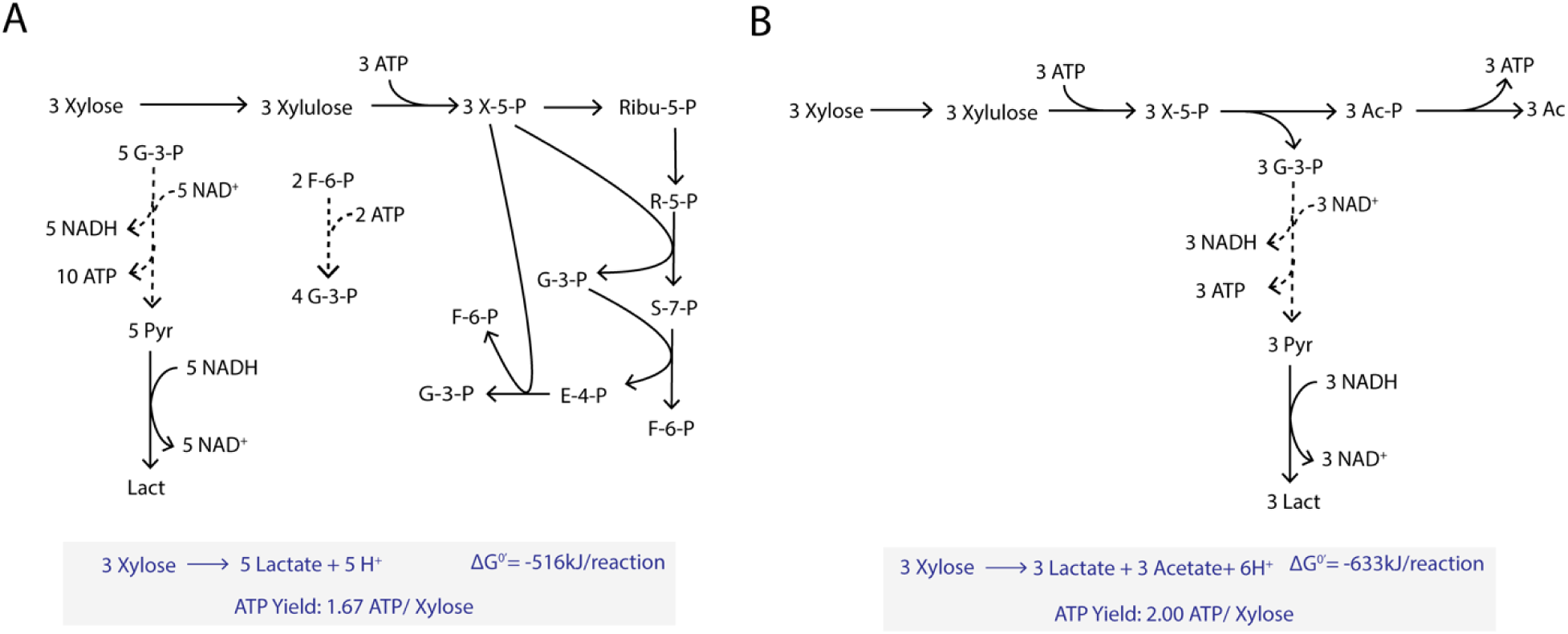
Comparison of ATP yields and thermodynamics with (A) consumption of xylose via the pentose phosphate pathway and (B) homolactic acid fermentation with the pentose phosphate pathway.

**Figure S6.**
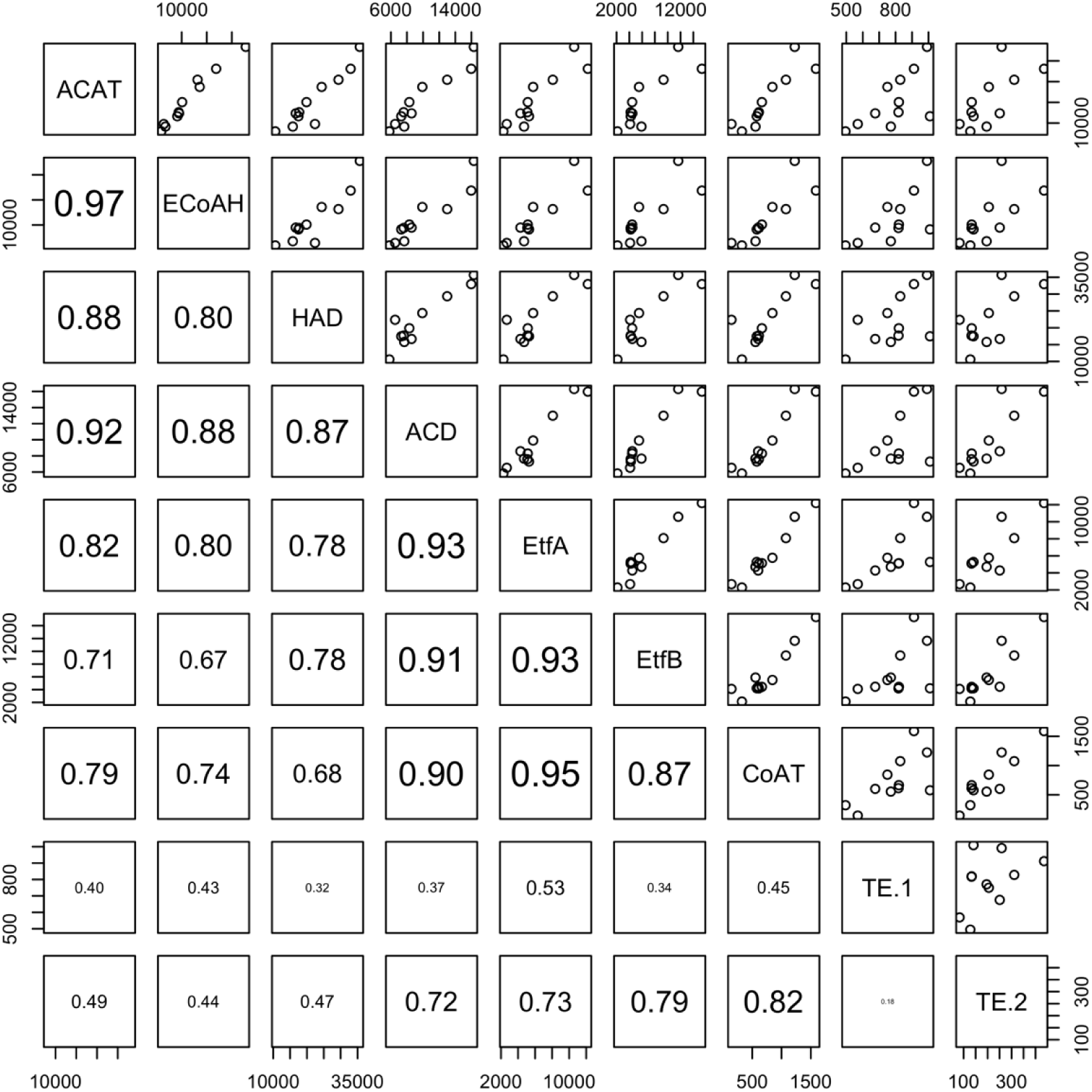
Pairwise comparison of transcript abundance for reverse β-oxidation genes in *Cand*. P. fermentans. Coefficients of determination are shown in bottom-left panels.

**Figure S7.**
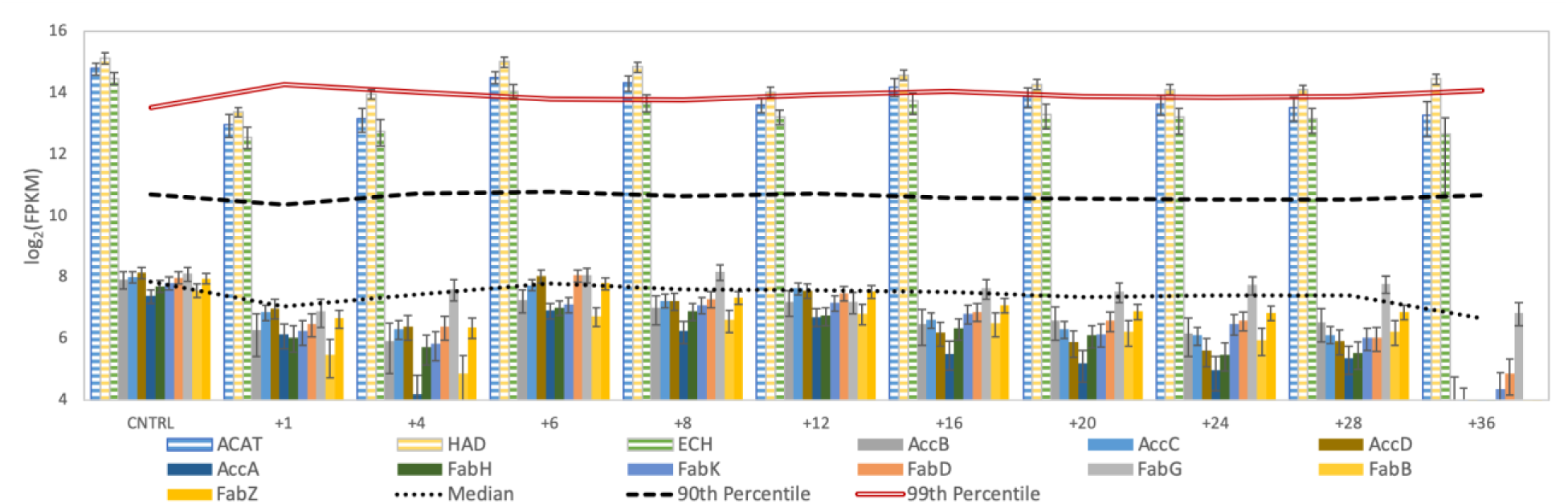
Abundance of reverse β-oxidation genes and fatty acid biosynthesis transcripts for *Cand*. P. fermentans. The genes are ordered as they are predicted to be ordered in the *Cand*. P. fermentans genome (Fig. 3J).

## LIST OF SUPPLEMENTARY DATA FILES

**Supplementary Data File 1:** Summary of DNA and RNA read mapping results

**Supplementary Data File 2:** Comparison of functional genome annotations for *Cand*. Weimerbacter bifidus (LCO1.1) with related genomes

**Supplementary Data File 3:** Comparison of functional genome annotations for *Cand*. Pseudoramibacter fermentans (EUB1.1) with related genomes

**Supplementary Data File 4:** Gene expression data for *Cand*. Weimerbacter bifidus (LCO1.1).

**Supplementary Data File 5:** Gene expression data for *Cand*. Pseudoramibacter fermentans (EUB1.1)

**Supplementary Data File 6:** Other functional genome annotations used for this study

**Supplementary Data File 7:** Summary of BLAST results for predicted thioesterase enzymes in *Cand*. Weimerbacter bifidus and *Cand*. Pseudoramibacter fermentans

**Supplementary Data File 8:** Summary of BLAST results for a predicted lactate dehydrogenase enzyme in *Cand*. Pseudoramibacter fermentans

